# Satb2 acts as a gatekeeper for major developmental transitions during early vertebrate embryogenesis

**DOI:** 10.1101/2020.11.23.394171

**Authors:** Saurabh J. Pradhan, Puli Chandramouli Reddy, Michael Smutny, Ankita Sharma, Keisuke Sako, Meghana S. Oak, Rini Shah, Mrinmoy Pal, Ojas Deshpande, Yin Tang, Rakesh Mishra, Girish Deshpande, Antonio J. Giraldez, Mahendra Sonawane, Carl-Philipp Heisenberg, Sanjeev Galande

## Abstract

Zygotic genome activation (ZGA) initiates regionalized transcription responsible for the acquisition of distinct cellular identities. ZGA is dependent upon dynamic chromatin architecture sculpted by conserved DNA-binding proteins. However, whether the tissue-specific transcription is mechanistically linked with the onset of ZGA is unknown. Here, we have addressed the involvement of chromatin organizer SATB2 in orchestrating these processes during vertebrate embryogenesis. Integrative analysis of transcriptome, genome-wide occupancy and chromatin accessibility revealed contrasting molecular functions of maternal and zygotic pools of Satb2. Maternal Satb2 represses zygotic genes by influencing the interplay between the pluripotency factors. By contrast, zygotic Satb2 activates transcription of the same group of genes during neural crest development and organogenesis. Comparative analysis of maternal versus zygotic function of Satb2 underscores how these antithetical activities are temporally coordinated and functionally implemented. We discuss the evolutionary implications of the biphasic and bimodal regulation of landmark developmental transitions by a single determinant.

---

The Cambrian explosion resulted in major diversification in the body plans of organisms^1^. Such diversification includes multiple sequentially progressive steps involving cell type specification, generation of distinct tissues, and organogenesis. Zygotic genome activation (ZGA) is the earliest developmental transition that imparts transcriptional competence to individual cells by incorporating positional information provided by localized patterning determinants^2^. ZGA is a conserved feature of early embryogenesis irrespective of the reproductive and morphological diversity. While ZGA relies upon genome-wide chromatin remodeling to establish transcriptional competence, how these genome-wide changes are translated into tissue-specific expression patterns remains largely unclear. Previous studies have identified specific regulators of ZGA and the precise mechanisms that control their mode of action to engineer regionalized gene expression have been investigated over the past decade^3–6^.

Spatiotemporal gene expression patterns are achieved by employing higher-order Gene Regulatory Networks (GRNs)^7^. GRNs are established via incorporation of novel gene families either de novo or by gene duplication and diversification leading to acquisition of novel functions^7^. An Important protein family which exhibits co-evolution and divergence consists of the special AT-rich binding proteins, SATB1 and SATB2. Both these proteins show high structural similarity and harbor a N-terminal ubiquitin-like domain (ULD), two DNA-binding CUT domains and a C-terminal homeobox domain^8^. Interestingly, SATB2 performs unique functions as a chromatin organizer during embryonic development, tissue specification and morphogenesis in vertebrates^9, 10^. SATB2 binds to multiple regulatory sites in a genome-wide manner and influences chromatin architecture thereby modulating target gene expression^11, 12^. In humans and mice, mutations in the *SATB2* locus result in cleft palate, craniofacial defects, developmental delay, cognitive disabilities and behavioral abnormalities^13–17^. SATB2 proteins exhibit a high level of conservation across vertebrates including zebrafish^18^. *satb2* is maternally deposited and ubiquitously expressed during early stages of zebrafish embryogenesis. Compromising Satb2 function using morpholinos resulted in severe developmental arrest^19^. Of note, later during organogenesis, loss of BMP and SHH signaling affect *satb2* expression in mandibles and pharyngeal arches of mouse and zebrafish embryos respectively^10, 20^. However, the molecular circuitry deployed by Satb2 to achieve early cell fate specification leading to primordial organ formation remains unclear.

Towards this, we generated a loss of function mutation in the zebrafish *satb2* locus. The mutation faithfully mimics important features of SATB2-associated syndrome in humans including craniofacial abnormalities which often arise from aberrant neural crest (NC) specification and migration^21, 22^. Neural crest cells (NCCs) are specialized multipotent cells that contribute to the development of cartilage, pigment cells, bones and connective tissue^23^. Transcriptome profiling of *satb2* mutants revealed global deregulation of genes involved in neural crest specification and migration. Specifically, our data demonstrate that zygotic Satb2 promotes expression of the positive regulators underlying NCC specification in zebrafish. Moreover, Chromatin accessibility analysis using Assay for Transposase-Accessible Chromatin with high throughput sequencing (ATAC-seq) performed using *satb2* mutant embryos at 14 somite stage suggested changes in global chromatin architecture via remodeling and nucleosome positioning. Genome-wide occupancy analysis of Satb2 in zebrafish and mice highlighted the conserved regulatory mechanisms across evolutionary scale. Surprisingly, our analysis revealed an unexpected novel function for maternally deposited Satb2. We demonstrate that unlike its zygotic counterpart, the maternal pool of Satb2 functions as a transcriptional repressor and controls the timing of ZGA by differentially regulating pluripotency factors. Stage-dependent transcription factor binding site (TFBS) prediction using genome-wide occupancy analysis has provided insights into mechanisms underlying the temporal activity of Satb2. Collectively, our data demonstrate that Satb2 functions in a versatile manner to influence crucial developmental transitions. Furthermore, its ability to participate in different biological contexts likely depends on diverse protein interactions and dynamic repertoire of genomic targets.

## Results

### SATB family proteins co-evolved with jawed fish

To gain a detailed insight into possible structural conservation among the SATB family proteins, we extracted the homologous sequence stretches by comparing the individual protein sequences. Initially SATB family proteins were thought to be restricted to the vertebrate lineage^24^. Subsequently, a distantly related ortholog *Defective proventriculus (Dve)* was identified both in fruit flies and roundworms. Interestingly, our phylogenetic analysis revealed the presence of SATB homologs in two other invertebrates; *Parasteatoda tepidariorum* (spider) and *Mizuhopecten yessoensis* (a molluscan-scallop) which include all three functional domains (Fig. 1a and Supplementary Fig. 1a). While Agnatha, a class of vertebrate jawless fishes, exhibits a single form of SATB protein, the presence of both SATB1 and SATB2 is only apparent in jawed fish and later species, suggesting these proteins presumably co-evolved with the evolution of the jaw structure. Importantly, based on domain architecture and phylogenetic affinities, SATB2 is closely related to the invertebrate homologs and hence can be regarded as the most ancient member of the family. Here, we focused on dissecting the function of SATB2 during evolution of developmental transitions underlying body plan determination.

**Fig. 1:**
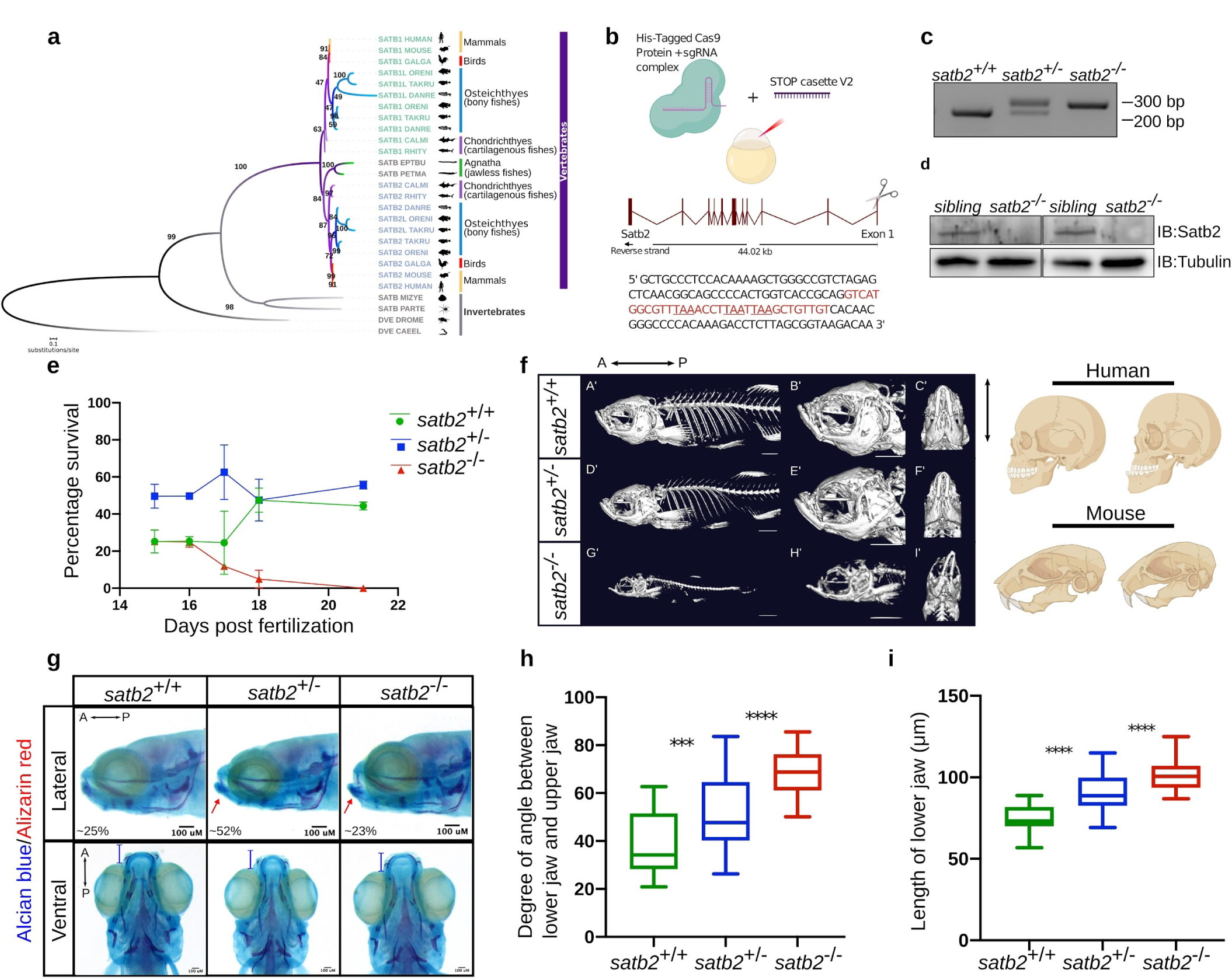
Loss of function of Satb2 in zebrafish leads to craniofacial defects and phenocopy function of mammalian homologues. **a**, Phylogenetic tree depicting evolution and divergence of SATB family proteins. SATB1 homologues are highlighted in green, SATB2 in blue whereas ancestors of SATB proteins are in grey. Organisms belonging to different classes are highlighted with different colours. Numbers on branching points represent bootstrap values. Organism labels; HUMAN- *Homo sapiens*, MOUSE- *Mus musculus*, GALGAL- *Gallus gallus*, ORENI- *Oreochromis niloticus*, TAKRU- *Takifugu rubripes*, DANRE- *Danio rerio*, CALMI- *Callorhinchus milii*, RHITY- *Rhincodon typus*, EPTBU- *Eptatretus burger*, PETMA- *Petromyzon marinus*, MIZYE- *Mizuhopecten yessoensis*, PARTE- *Parasteatoda tepidariorum*, DROME- *Drosophila melanogaster* and CAEEL- *Caenorhabditis elegans*. Additional copies of SATB1 and SATB2 were denoted with SATB1L (SATB1-like) and SATB2L (SATB2-like). **b**, Schematic of CRISPR-cas9 mediated mutant generation by introducing the STOP cassette (highlighted in red) in exon1 of the *satb2* gene resulting in loss of function mutant allele. **c**, Identification of mutant allele by genotyping (60 bp insertion) and **d**, confirmation of loss of Satb2 between siblings and homozygous larvae at 48 hpf by immunoblot. Tubulin was used as a loading control. **e**, Lifespan analysis of zygotic *satb2* mutants depicted by a line graph. Wild-type (green), Heterozygous (blue) and Homozygous mutants (Red). Error bar represents +/− S.D. of two independent experiments with n=48. **f**, Micro CT images of adult zebrafish (4 months old) to visualize defects in skeletal and craniofacial structures. Lateral view A’, D’, G’, magnified lateral view B’, E’, H’ and ventral view C’, F’, I’ of Wild-Type, Heterozygous and Mutant for Satb2 respectively are shown. Ventral view images are not scaled to attain a maximum field of view highlighting detailed structural deformities independent of the size of fish. Image is a representative of N=5. Corresponding schematics depicting similarity with *SATB2* mutation reported in humans and mice. **g**, Alcian blue/Alizarin red staining to visualize craniofacial defects at early larval stages, 15 dpf. Abnormal jaw protrusion in heterozygous and homozygous mutants is indicated by a red arrow. Numbers in respective boxes signifies the percentage of the larvae from the total population showing class of phenotype and genotype correlation. Degrees of the angle between the lower jaw and upper jaw **h**, and a total length of protrusion measured from ventral view **i**, as indicated in panel F, are plotted using a box plot. *** indicates p-value of significance 0.001 and **** indicates p-value of significance 0.0001 as determined by student’s two-tailed t-test.

### Disruption of Satb2 function leads to defective craniofacial patterning

In zebrafish embryos, compromising Satb2 activity using morpholinos resulted in severe developmental defects including epiboly arrest. These phenotypes were attributed to aberrant exocytosis and endocytosis^19^. We reasoned that the pleiotropic phenotype could be a result of depletion of the maternal as well as zygotic pool of Satb2. To uncouple the maternal versus zygotic requirement of Satb2, we introduced a mutation in *satb2* locus using the CRISPR-cas9 system^25^ by inserting a STOP cassette into the first exon, leading to premature termination (Fig. 1b and Supplementary Fig. 1c). The absence of Satb2 in the mutant fish was confirmed by immunoblot analysis (Fig. 1c,d).

Homozygous zygotic *satb2* mutants showed poor survival to adulthood with most of the *satb2*^-/-^ fish being depleted from the pool after 16 days post fertilization (Fig. 1e). However, we could occasionally recover homozygous survivors (∼1%) from the total population (n=400). Such survivors displayed severe deformities and reduced body size compared to their heterozygous siblings (Supplementary Fig. 1b,c). The homozygous fish were also sterile. Given the requirement of Satb2 during craniofacial development and bone formation in mammals, we recorded CT scans of homozygous survivors, heterozygous mutant siblings, and wild-type fish (n=5). *satb2* mutant fish displayed deformed mandibles with a high degree of lower jaw protrusion, reduced maxilla with larger eye pockets and abnormal skull size and shape (Fig. 1f). Supporting the haploinsufficient nature of Satb2 function, heterozygous fish also showed less severe yet detectable cranial structural abnormalities. Thus, zebrafish *satb2* mutant mimics the craniofacial defects observed in humans and mice^14^ (Fig. 1f).

To trace back the onset of these defects during early development, we performed alcian blue: alizarin red staining on larvae obtained by mating *satb2* heterozygous adults. Larvae were collected 15 days post-fertilization, a day prior to depletion of zygotic *satb2^-/-^* fish from the pool. As in the case of adult survivors, we observed craniofacial defects in the homozygous larval samples (Fig. 1g). We also detected an increase in the angle between the mandibles and maxilla in heterozygous and homozygous mutant larvae as compared to wild-type siblings (Fig. 1h). Measurement of the total length of the lower jaw confirmed significantly higher protrusion in zygotic s*atb2* mutants (Fig. 1i). Thus, our data demonstrate that zygotic loss of Satb2 in zebrafish leads to defective cranial and skeletal morphogenesis providing us with a unique tool to understand its function during initial events of vertebrate organogenesis.

### Satb2 potentiates early neurogenesis and neural crest specification

Aberrant neural crest specification and migration have been correlated with severe craniofacial defects^21, 22^. In the developing larvae, *satb2* is strongly expressed in the pharyngeal arches, a reservoir of NCCs^18^. This prompted us to investigate the possible involvement of Satb2 in NC development. We selected three developmental time points that correspond to specific stages of NC differentiation. To study NC induction, NC specification and differentiation we analyzed gastrulating embryos at 80% epiboly, embryos at 5-6 somites stage (ss) and 14 ss respectively^26^. We performed gene expression analysis in wild-type siblings and Satb2^-/-^ embryos at the corresponding stages (Fig. 2a). PCA plots and Pearson correlation analysis for all stages showed significant reproducibility between the replicates (Fig. 2b and Supplementary Fig. 2c). Our analysis revealed that loss of Satb2 function resulted in deregulation of numerous genes across all developmental stages (658 at 80% epi, 661 at 6 ss, 3015 at 14 ss). Both at the level of gene expression and in terms of number of deregulated genes the phenotype was most drastic at 14 ss. This suggested that Satb2 plays an essential role during NC differentiation (Figures 2B and S2C). Interestingly, we observed minimal overlap between differentially expressed genes across developmental stages (Figure 2C). Moreover, at 14 ss, the deregulated targets included known molecular components of NC development such as *sox10*, *sox9a*, *pax3a* and *zic1*^27–31^ (Fig. 2d).

**Fig. 2:**
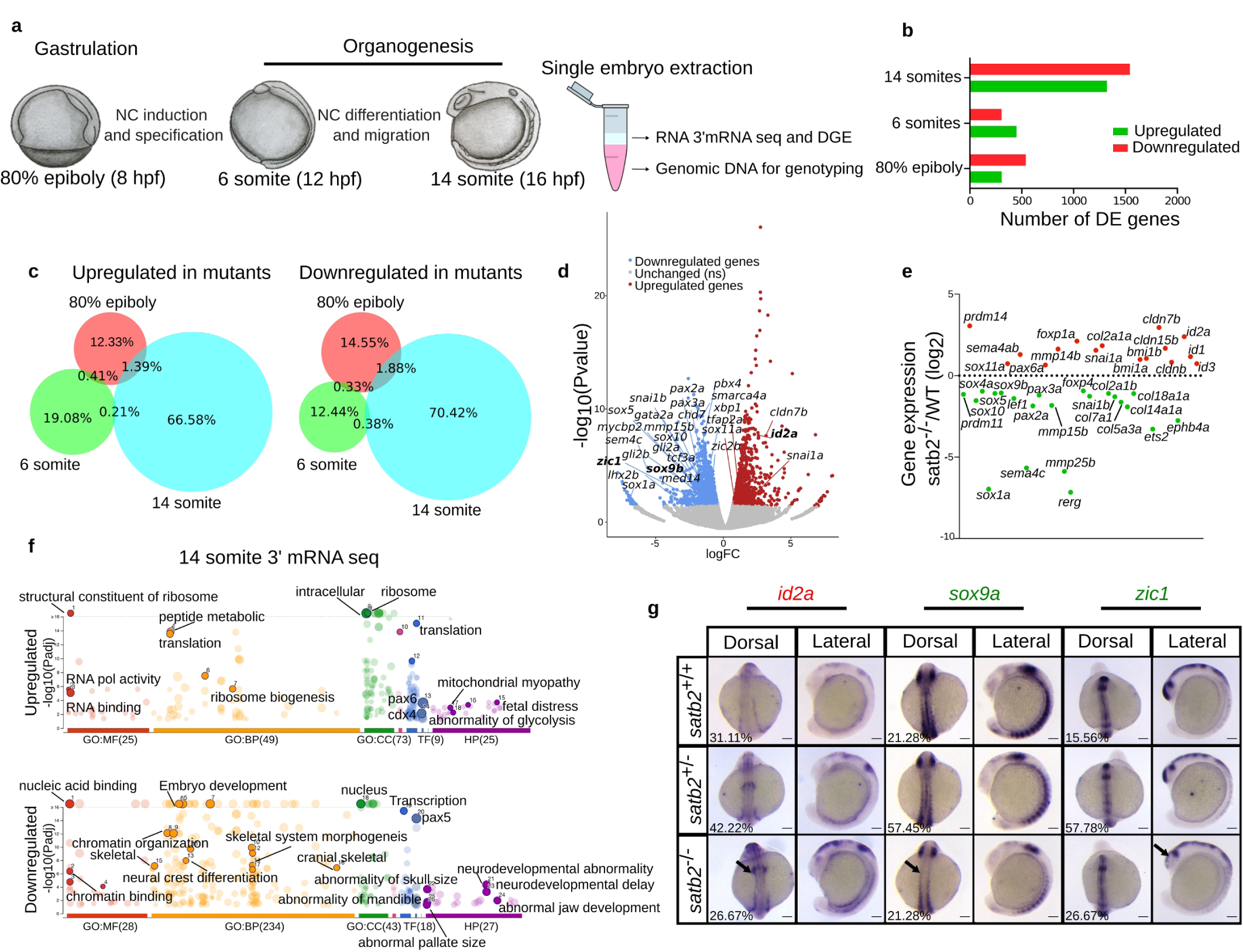
Deregulation of gene expression in *satb2* mutants at the onset of organogenesis leads to defective neurogenesis and craniofacial patterning. **a**, Experimental strategy for gene expression analysis using 3’mRNA sequencing at 80% epiboly, 6 ss and 14 ss. **b**, Bar plot for the number of upregulated (red) and downregulated (green) genes at respective stages. **c**, Venn diagram showing percentage overlap between differentially expressed genes in *satb2^-/^*^-^ at different stages. **d**, Volcano plot of differentially expressed genes at 14 ss embryonic stage. DE genes with FDR < 0.1 and log2 Fold change > +/− 0.58 are coloured as red for upregulated, blue for downregulated and grey for unchanged or non-significant genes. Few significantly differentially expressed genes representing GO categories neural crest differentiation, neurogenesis and cranial skeletal system development are highlighted. **e**, Scatter plot for Log_2_ Fold change in gene expression of various gene families belonging to the above-mentioned gene ontology categories **f**, GO analysis of dysregulated genes at 14 ss highlighting significant classes under molecular function (MF), biological processes (BP), Cellular components (CC), Transcription factors (TF) and Human pathology (HP) categories. **g**, Dorsal and lateral view of WISH for *Id2a* (upregulated), *sox9a* and *zic1* (downregulated). Arrows in the respective images mark the region of maximum differences in expression pattern.

Consistent with the selective nature of Satb2’s function, many paralogs within the gene families such as *prdm (prdm11, prdm14)*, *foxp (foxp4*, *foxp1a)* and *snai (snai1 and snai1b)* were differentially regulated by Satb2 (Fig. 2e). Gene ontology analysis of upregulated genes at 14 ss showed significant enrichment in pathways involved in RNA polymerase activity, ribosome biogenesis and mRNA translation. Importantly, downregulated gene sets were enriched for the components of skeletal morphogenesis, NC differentiation and early embryonic developmental pathways. Supporting a conserved function of zebrafish Satb2, mutations in these target genes have been correlated with neurodevelopmental and craniofacial abnormalities in humans (Fig. 2f). To validate these changes, we performed whole-mount in situ analysis for *id2a*, *sox9a* and *zic1*. *satb2^-/-^* mutants exhibited reduced expression of *sox9a* in the neural tube and *zic1* in the developing head. In contrast, *id2a*, a negative regulator of NCC specification, was considerably upregulated (Fig. 2g). Collectively, these findings show that Satb2 is required during initial events of NC development and loss of Satb2 function results in perturbation of the underlying gene expression programs.

### Genome-wide occupancy analysis confirms the direct regulation of downstream target genes by Satb2

To investigate the molecular basis of Satb2 function, we performed chromatin immunoprecipitation followed by high throughput sequencing (ChIP-seq) for the corresponding stages of embryonic development. To this, we generated a polyclonal antibody against a peptide spanning the N-terminal region of Satb2 (see methods). To test the antibody, we overexpressed 3xFLAG-Satb2 from one-cell stage and performed ChIP-seq at the dome stage using either anti-Flag or anti-Satb2 antibodies. Confirming the specific nature of the antibody, we observed a very high correlation (Pearson correlation 0.80) between ChIP signal intensities across these two conditions (Supplementary Fig. 3b).

Next, we performed ChIP-sequencing in replicates using anti-Satb2 antibody and observed a very high correlation (Supplementary Fig. 3c). To identify Satb2 binding sites in open chromatin regions (OCR), we intersected consensus peaks with Ensembl of chromatin accessibility across various developmental stages (GSE106428, GSE130944, GSE101779) yielding 27420 binding sites at 80% epiboly, 36,444 sites at 6 ss and 22,333 sites at 14 ss respectively. Genomic distribution analysis suggested that Satb2 occupies both the promoter and non-promoter regions (intergenic + intronic + exonic + TTS) (Fig. 3a). Interestingly, this data indicates that genes involved in early neurogenesis and NC development such as *sox10*, *sox9a*, *zic1* and *zic4* are potentially book-marked by the presence of Satb2 as early as at 80% epiboly stage (Fig. 3b).

**Fig. 3:**
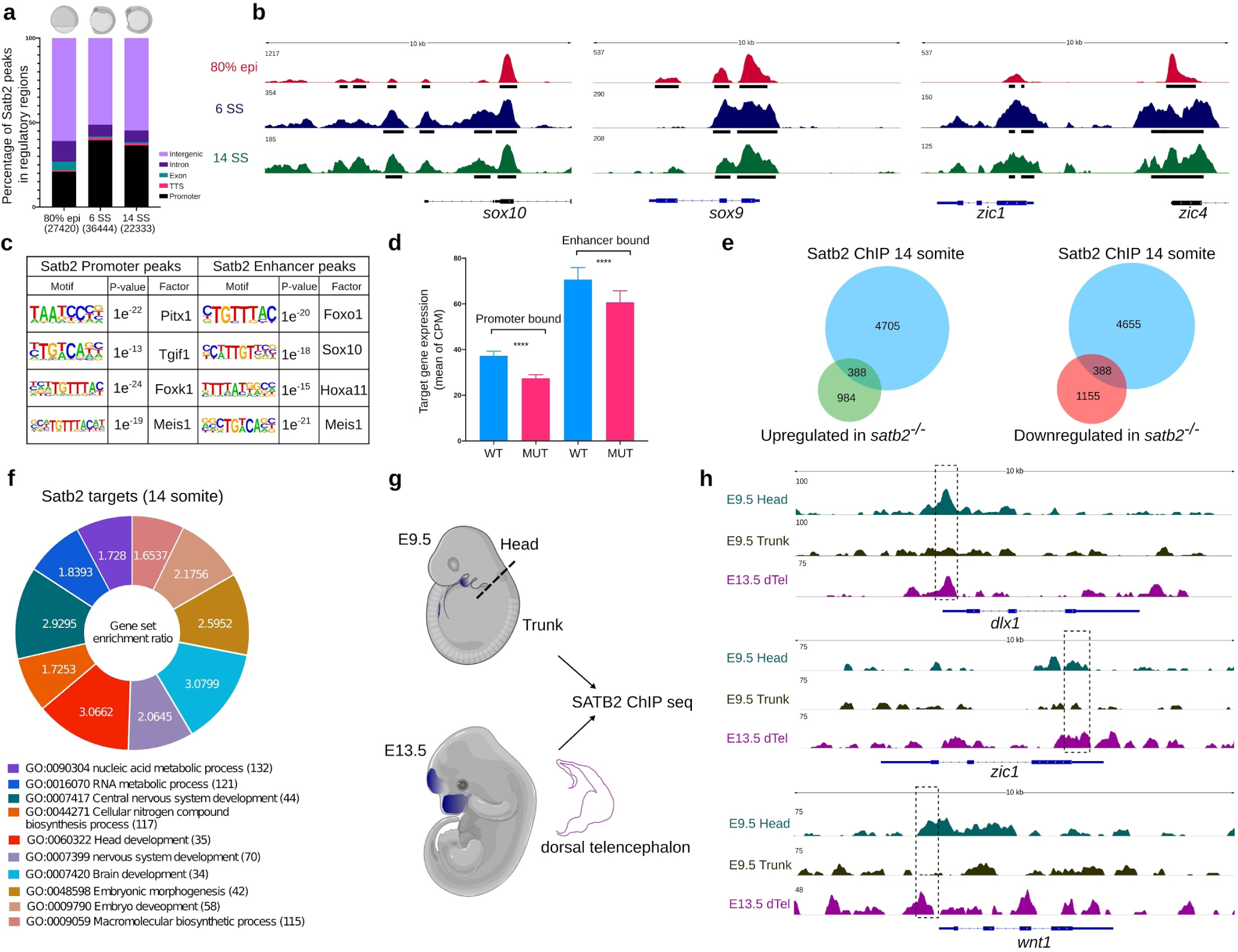
Genome-wide occupancy analysis of Satb2 underscores the molecular mechanism underlying Satb2 function. **a**, Genome-wide distribution pattern of Satb2 occupancy in the regulatory regions. percentage of peaks overlapping with OCR at 80% epiboly, 6 ss and 14 ss. Numbers in the brackets indicate the total number of Satb2 peaks at the corresponding developmental stage. **b**, Integrative genomics viewer (IGV) snapshot of Satb2 occupancy on genomic loci of *sox10*, *sox9a* and *zic1*. Black coloured blocks represent enriched regions determined by the peak calling algorithms. **c**, Motif enrichment analysis for Satb2 binding sites at the promoter (+/− 2 Kb) and enhancer regions (overlapping with H3K4me1 binding sites) matched to known motif sets of danRer10 genome assembly. **d**, Bar plots indicating normalized gene expression (Counts Per Million) of promoter- and enhancer-bound genes in wild type and Satb2 ^-/-^ mutants at 14 ss. **e**, Venn analysis depicting the overlap between Satb2 bound genes and differentially regulated genes at 14 somites in the Satb2^-/-^ mutant. **f**, GO analysis for biological processes of Satb2 occupied genes at 14 ss. Numbers for each category of doughnut plot represents enrichment ratio, FDR < 0.05. **g**, Schematic of dissections performed for the ChIP-seq experiment in mouse embryos at E9.5 (head and Trunk) and E13.5 (dorsal telencephalon). Dotted lines represent the site of the incision. **h**, IGV snapshot of mouse SATB2 occupancy profile on the genomic loci of *dlx1*, *zic1* and *wnt1.* Dashed line boxes highlight enriched regions in E9.5 head and E13.5 dTel which are absent in the E9.5 trunk region.

To assess whether Satb2 binds to putative enhancer regions of these genes, we intersected the Satb2 regulatory peak set with previously published dataset for poised enhancers across developmental stages (GSE32483, GSE74231) as marked by H3K4me1^32^. Satb2 peaks from the promoter and putative enhancer regions were further subjected to motif analysis. Interestingly, we observed a strong enrichment for transcription factors including Pitx1, Tgif1, Foxk1 and Meis1 at the promoters (Fig. 3c)). Mutations in these genes also affect craniofacial development in mice and humans^33–37^. Moreover, enhancer bound regions showed strong enrichment for binding sites of known regulators of NCC development such as Foxo1, Sox10 and Hoxa11^27, 28, 38, 39^. To assess whether binding of Satb2 influences gene expression, we compared corresponding transcript levels between wild-type and *satb2* mutants. Indeed, genes containing Satb2 binding sites within promoters and/or enhancers were significantly down-regulated in the mutants (Fig. 3d).

We also performed reciprocal analysis comparing transcript levels between wild-type and mutants obtained from RNAseq experiments and correlated it with Satb2 occupancy. To achieve this, we annotated Satb2 bound regions to the nearest genes and analyzed the overlap between the two. This analysis demonstrated that Satb2 occupied both upregulated and downregulated sets of genes (Fig. 3e). Thus, we concluded that Satb2 exhibits bimodal activity through directly binding to cis-regulatory sequences. To confirm this notion, we subjected Satb2 genomic targets to gene ontology analysis. Consistent with our earlier observation, upregulated gene set showed enrichment for biological processes involved in nucleic acid metabolism, RNA metabolism and nitrogen biosynthesis; whereas downregulated gene set was enriched for processes involved in central nervous system development and morphogenesis (Fig. 3f). Taken together, the correlation analysis between Satb2 occupancy and mutant transcriptome underscores a distinct function of zygotic Satb2 in regulating early neurogenesis and RNA metabolism.

### Direct binding to cis-regulatory elements is a conserved function of Satb2

To extend these observations, we decided to perform ChIP-seq analysis on developing mouse embryos. Use of developing mouse embryos provided us with a window to study spatial regulation by Satb2 during NC specification. At E9.5 we dissected the head (source for cranial NCCs) and the trunk tissue (source of NCCs contributing to the peripheral nervous system). As a positive control for ChIP experiments, we isolated the dorsal telencephalon (dTel) at E13.5 to enrich the population expressing Satb2 in the developing cortex (Fig. 3g). The ChIP-seq analysis in mice indeed showed significant enrichment for Satb2 on the regulatory regions of bonafide NCC markers such as *dlx1*, *zic1* and *wnt1* specifically in the head tissue. However, Satb2 occupancy was not enhanced in trunk tissue for these regulatory regions, suggesting locus-specific and spatially restricted regulation by mouse SATB2 (Fig. 3h). Taken together, our data suggests that functional regulation by Satb2 through direct binding at cis-regulatory elements of target genes is a conserved mechanism across vertebrates.

### Loss of Satb2 results in reduced chromatin accessibility and perturbed nucleosome positioning

As Satb2 activity requires binding to target DNA sequences, we were interested in examining if Satb2 also modulates chromatin architecture. To assess this, we performed ATAC-seq in wild-type siblings and Satb2^-/-^ mutants at 14 ss. Satb2^-/-^ mutants yielded a significantly lower number of OCRs. Interestingly, the percentage of OCRs at intergenic loci dropped significantly as compared to the promoters (Fig. 4a). Clustering (K-means=4) of ATAC-seq signals around the +/− 2 Kb centre of Satb2 peaks showed significant reduction across the genome in *satb2* mutants (Fig. 4b and Supplementary Fig. 4a). The gene ontology analysis revealed cluster 2 to be enriched for biological processes such as somite formation, embryonic morphogenesis, and craniofacial development (Fig. 4c) whereas, Cluster 1 showed enrichment for various metabolic processes (Supplementary Fig. 4b). Motif enrichment analysis of cluster 2 yielded significant overlap with binding sites for known regulators of neurogenesis and NCC specification such as Sox3, Pitx1 and Sox10 (Fig. 4d). To identify differential chromatin accessibility regions, we subjected the ATAC-seq data to DiffBind analysis. We found a high number of regions (6861) with loss of chromatin accessibility in mutants as compared to only 141 regions with a gain of accessibility (Fig. 4e). Furthermore, we observed that modulation of chromatin accessibility primarily influences intergenic regions of genes including those involved in NCC specification (Fig. 4a,f,g).

**Fig. 4:**
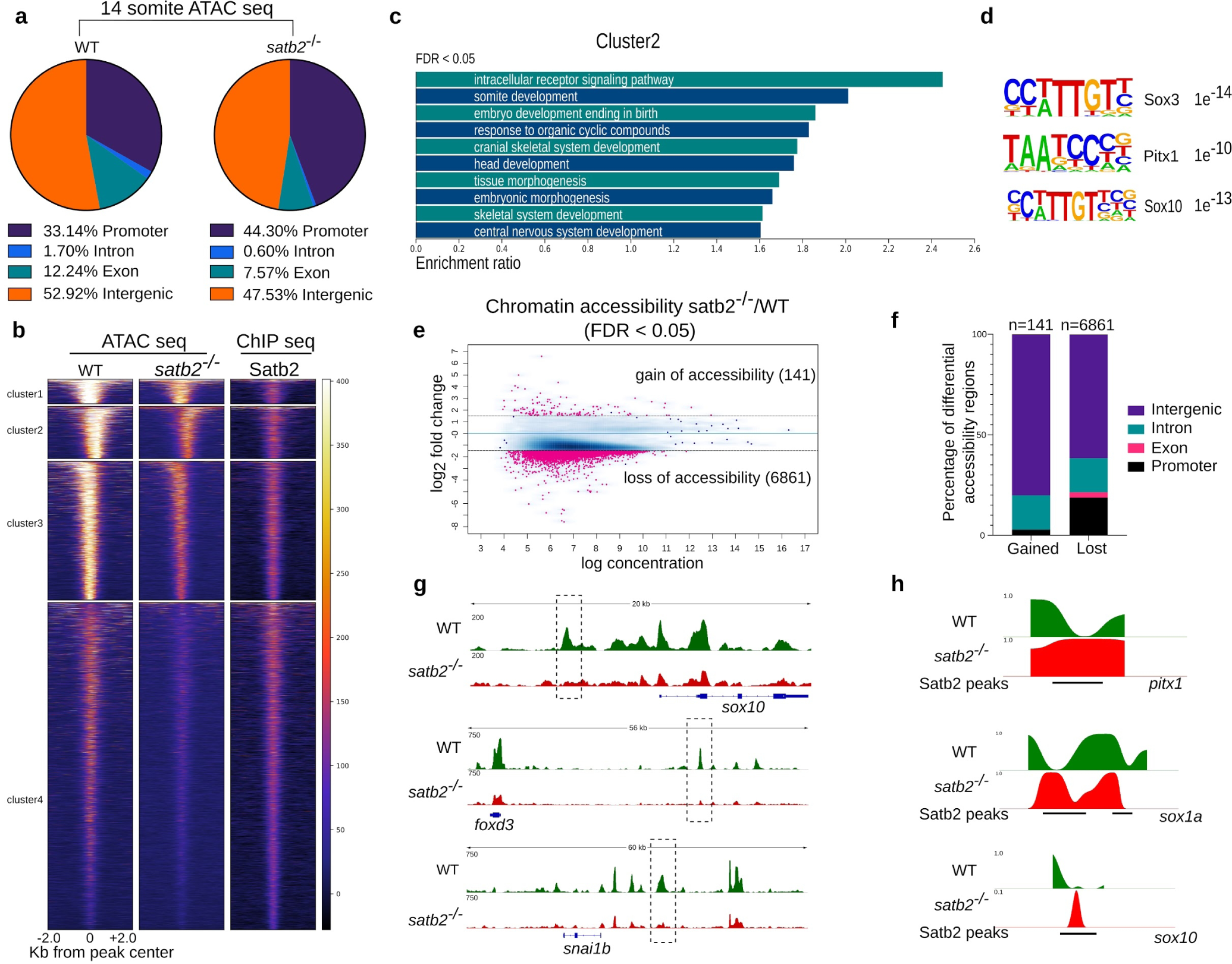
Loss of Satb2 affects chromatin accessibility. **a**, Genomic distribution of ATAC-seq peaks in wild type and Satb2^-/-^ mutant at 14 ss as a percentage of the total. **b**, Heatmaps for clustering (K-means 4) of chromatin accessibility around +/− 2 Kb of Satb2 ChIP-seq peak regions. Colorbar represents the degree of chromatin accessibility from low (blue) to high (white). **c**, GO analysis of cluster 2 highlighting biological processes involved in neural and craniofacial development. **d**, Enriched TFBS are represented for cluster 2 **e**, Differential chromatin accessibility between wild type and Satb2^-/-^ mutant embryos represented as binding affinity (FDR < 0.05). Y-axis represents Log2 fold change. Regions with Log2 fold (> +/− 0.58) were considered significant and marked by solid lines. **f**, Genome-wide distribution of differentially gained or lost regions in Satb2^-/-^ mutant embryos represented as a percentage. **g**, IGV snapshot of ATAC-seq peaks over the genomic loci for enhancer regions of neural crest markers *sox10*, *foxd3* and *snai1b*. Dashed line boxes highlight regions which are identified as differentially expressed through Diffbind analysis. **h**, Nucleosome occupancy profile calculated using ATAC-seq showing changes in nucleosome phasing at Satb2 regulatory sites for *pitx1*, *sox1a* and *sox10*.

Reduced chromatin accessibility is a critical determinant of gene regulation. However, genome organizer proteins can directly affect nucleosome positioning^40^. To test whether Satb2 has any impact on nucleosome positioning, we estimated nucleosome occupancy from ATAC-seq data using the nucleoATAC suit^41^. To our surprise, we observed reduced occupancy signals in mutants compared to wild-type at the transcription start site (TSS) as well as at the Satb2 binding regions (Supplementary Fig. 4c,d). To analyze this further, we compared the nucleosome distribution at specific Satb2 target gene loci. We observed disruption in nucleosome phasing which resulted in fuzzy nucleosome patterns, particularly at the Satb2 binding sites (Fig. 4h and Supplementary Fig. 4e). Of note, we did not observe such change in nucleosome positioning at the TSS for genes which are upregulated in Satb2^-/-^ mutants (Supplementary Fig. 4c). These findings suggest that down-regulation of Satb2 target genes could be resulting due to disruption in defined nucleosome phasing. In sum, the chromatin accessibility analysis uncovered multiple mechanisms through which Satb2 actively ensures sustained gene expression to drive neurogenesis and NCC development.

### Tracing back the activity of Satb2 during zygotic genome activation

Data presented in the previous sections demonstrate a direct regulatory role of zygotic Satb2 during early neurogenesis and NCC development. Intriguingly, Satb2 is also maternally deposited and is thought to function primarily during epiboly^19^. Analysis of publicly available time-series transcriptome data during zebrafish embryogenesis^42^ revealed bi-phasic expression of *satb2*. Maternal *satb2* is significantly reduced at the onset of ZGA and maintained at a minimal level before zygotic *satb2* starts expressing during organogenesis (Fig. 5a).

**Fig. 5:**
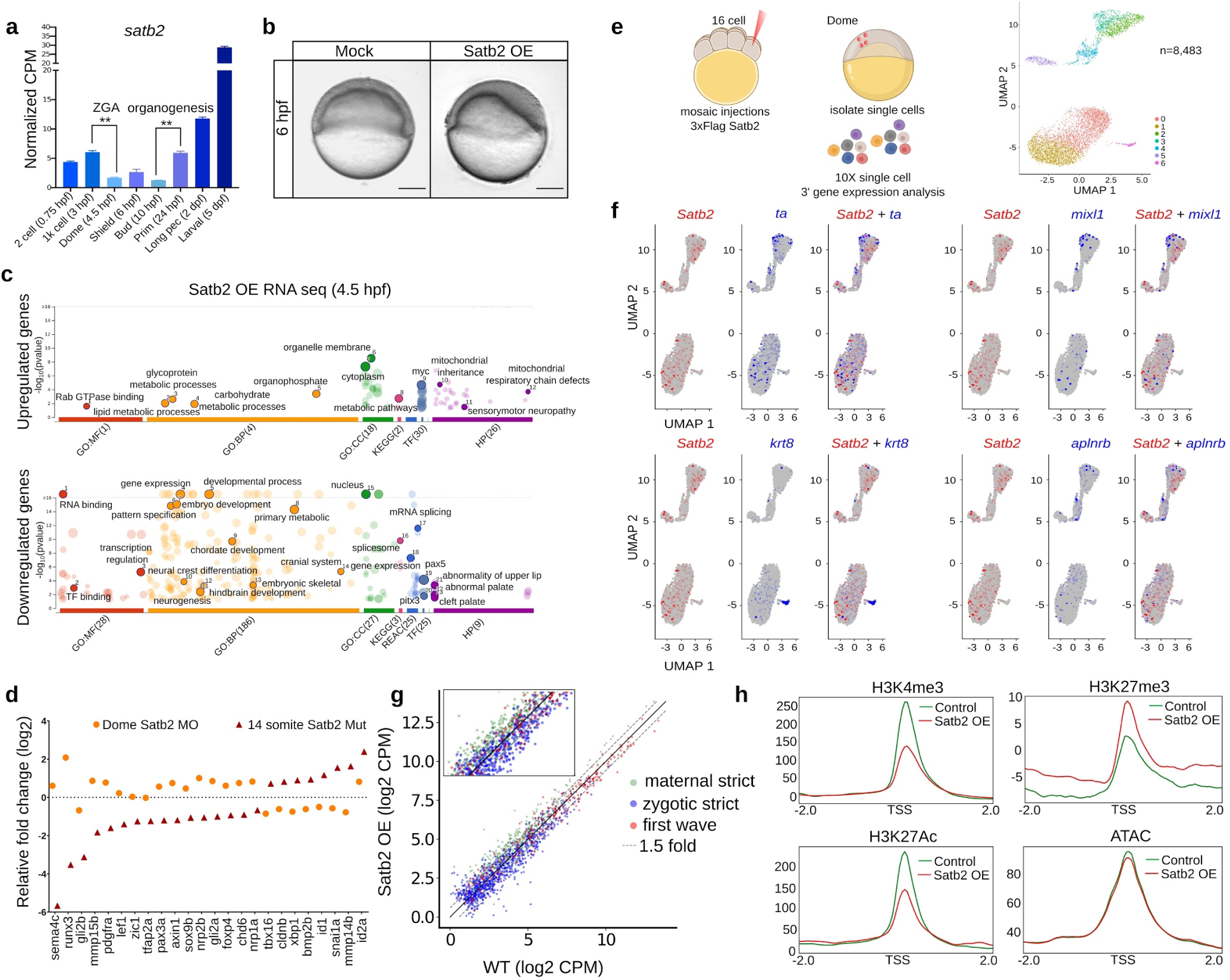
Maternal Satb2 functions as a transcriptional repressor prior to ZGA. **a**, Stage-specific gene expression analysis highlighting bi-phasic expression of Satb2 during MZT and organogenesis. Normalized CPM values for each stage are used for the analysis. (** represents P-value of < 0.05 as measured by students two-tailed t-test) **b**, Lateral view of zebrafish embryos at 6 hpf injected with 200 pg eGFP (control) and 3xFlag-Satb2 mRNA. **c**, GO analysis for differentially expressed genes upon over-expression of Satb2 at 4.5 hpf. Significant ontologies under molecular function (MF), biological processes (BP), Cellular components (CC), Reactome (REAC), Transcription factors (TF) and Human pathology (HP) categories are highlighted. Numbers in brackets signify the total number of ontologies enriched under each category. **d**, Relative expression values for genes involved in neural crest differentiation upon morpholino mediated depletion of maternal Satb2 (yellow circles) analyzed at 4.5 hpf and in zygotic Satb2^-/-^ mutants (red triangles) analyzed at 14 ss, highlighting opposite effects on gene regulation by maternal and zygotic forms of Satb2 (FDR<0.1). **e**, Experimental strategy for single-cell analysis of embryos with mosaic overexpression of Satb2 at 4.5 hpf. UMAP clustering illustrates 8 different clusters based on gene expression patterns from 8,483 single cells. **f**, FeaturePlots for exemplifying negative correlation between cells expressing *satb2* (blue) and zygotic factors *ta*, *mixl1*, *krt8* and *aplnrb* (red). **g**, Biplot depicting the effect of overexpression of Satb2 on maternal (green), zygotic (blue) and first wave zygotic genes (red). Log_2_ CPM values are used for the analysis. The dotted line marks genes which show more than 1.5 fold differences in Log2 CPM values. Inset is a zoomed version to highlight the differences. **h**, Average mean density profiles around +/− 2 Kb region around TSS of ‘zygotic genes’ for H3K4me3 (activation mark), H3K27me3 (repressive mark), H3K27Ac ( Activation mark) and ATAC-seq (chromatin accessibility) upon Satb2 overexpression (red) compared to control (green).

To assess the function of maternally deposited *satb2*, we attempted to generate maternal-zygotic (MZ) mutants for *satb2*. As in the case of mice, zygotic *satb2* mutants show poor survival and the rare survivors are infertile^17^ (Fig. 1e). To circumvent this difficulty, we attempted germ line transplantations but resulting fish were also infertile (data not shown). Thus, we could not obtain MZ mutants to directly examine the function of the maternal pool of Satb2.

We next turned our attention to the biological relevance of the reduction of maternally deposited s*atb2* at the onset of a major wave of ZGA. To assess this, we overexpressed Satb2 from 1 cell stage (Supplementary Fig. 5a,b). Curiously, as in the case of morpholino-mediated knockdown, overexpression of Satb2 resulted in significant developmental defects^19^ (Fig. 5b). This observation suggested that temporally controlled reduction of Satb2 could be biologically relevant. To gain further insight into Satb2 function, in the following, we have carefully examined functional consequences of the sustained presence of Satb2 in early zebrafish embryos.

Satb2 overexpressing embryos fail to survive beyond 8 hpf indicating possible early defects in cell type specification and morphogenesis. Closer inspection of these embryos uncovered thickening of the dorsal organizer region. Similarly, In *Xenopus*, *satb2* expression was enriched in the dorsal organizer suggesting its involvement in dorsoventral patterning^43^.

Next, we performed transcriptome analysis of Satb2 overexpressing embryos. To rule out the effect of developmental delay on the gene expression we used embryos at 4.5 hpf (dome stage). As in the case of zygotic satb2 mutants, overexpression of Satb2 also resulted in significant changes in gene expression (Supplementary Fig. 5c). However, in contrast to zygotic loss of Satb2, gene ontology analysis revealed that upregulated genes primarily constitute part of metabolic processes whereas downregulated genes were enriched in embryonic development, pattern specification, skeletogenesis, neurogenesis and NC differentiation (Fig. 5c). These data argue that maternal Satb2 potentially acts as a repressor of the group of genes that are subsequently activated by zygotic Satb2 during organogenesis. (Figure S5D).

To confirm this directly, we inactivated satb2 function using two different morpholinos targeting the translation start site and splice junction resulting in early developmental arrest. (Supplementary Fig. 6a). Five base Mismatch morpholino was used as a control. Moreover, a morpholino-resistant version of satb2 mRNA could partially rescue the phenotype (Supplementary Fig. 6b). Importantly, in contrast to Satb2 overexpression, knockdown embryos exhibit reduced expression of dorsal marker *chrd* (Supplementary Fig. 6c).

Next, we performed bulk mRNA-seq upon Satb2 knockdown and specifically analyzed the expression of genes involved in neurogenesis and NC development which are positively regulated by zygotic Satb2. As anticipated, markers for neurogenesis and NC development were prematurely upregulated (Fig. 5d and Supplementary Fig. 6d). However, compromising maternal Satb2 levels did not affect exocytosis and endocytosis pathway components^19^. These results establish that unlike zygotic Satb2, the maternal pool of Satb2 is involved in transcriptional repression and possibly regulates the same group of genes in an opposite manner.

### Satb2 functions in a cell autonomous manner

To assess the nature of Satb2’s function during ZGA we employed single-cell gene expression analysis. We overexpressed Satb2 in a mosaic manner by injecting 3xFLAG-Satb2 in one of the cells at 16 cell stage embryos and harvested embryos at 4.5 hpf to prepare single-cell suspension (Fig. 5e). This strategy also enabled us to obtain wild-type cells from the same source. We generated a high-quality dataset for 3’ gene expression in 8483 single cells using scRNA-seq (Supplementary Fig. 5e,f). UMAP clustering of these cells identified 7 distinct clusters (cluster 0 to cluster 6) based on the correlation of gene expression patterns (Fig. 5e). Next, we analyzed the expression of individual genes as specific features and generated spatial maps for *satb2* and developmental genes which are found to be negatively regulated by Satb2 through our bulk mRNA-seq analysis. To exemplify, we represent spatial maps for *satb2*, *ta*, *mixl1*, *krt8* and *aplnrb* and highlight a negative correlation between the expression of *Satb2* and these developmental genes (Fig. 5f). In conclusion, mosaic overexpression followed by scRNAseq indicated that Satb2 functions in a cell-autonomous manner.

### Maternal Satb2 modulates local chromatin landscape

Lee et al. have previously characterized early developmental genes into strictly maternal, strictly zygotic and first-wave zygotic genes^4^. Upon overexpression of Satb2, genes belonging to strictly zygotic, and first wave zygotic genes were significantly downregulated (Fig. 5g and Supplementary Fig. 5g). By contrast, maternally deposited genes required to support early growth were upregulated^44^ (Fig. 5c,g).

To assess if repression of zygotic genes is correlated with corresponding epigenetic changes, we profiled occupancy of H3K4me3 (activation mark), H3K27me3 (repressive mark), H3K27Ac (activation mark) and analyzed chromatin accessibility (ATAC-seq) focusing on TSS of ‘zygotic strict’ genes. Curiously, we observed depletion and increase in the activation and repressive marks respectively despite a very minimal effect on chromatin accessibility (Fig. 5h). This could be presumably due to transient regulation of epigenetic landscape or active compensatory mechanisms.

### Interplay between Satb2 and pioneer factors at ZGA

Role of the pioneer factors such as Pou5f3 (homolog of OCT4), Nanog and SoxB1 has been well established in activating ZGA^4, 5^. To test, if maternal Satb2 is preventing precautious transcription of zygotic patterning genes by targeting pioneer factor(s) we investigated if these factors are specifically regulated by Satb2. Supporting the notion, RNAseq analysis and whole-mount *in situ* demonstrated downregulation of *pou5f3*. Notably, unlike *pou5f3*, *sox19b* was upregulated upon overexpression of Satb2, suggesting influence of Satb2 on the pioneer factors is qualitatively non-uniform (Fig. 6a, 6b). Interestingly, many of the direct targets of Pou5f3 and Sox2 showed significant overlap with negative targets of Satb2 (Fig. 6c, 6d). We subsequently confirmed that the negative regulation between Satb2 and Pou5f3 is not unique to zebrafish but is conserved across vertebrates (Fig. 6e and Supplementary Fig. 7a).

**Fig. 6:**
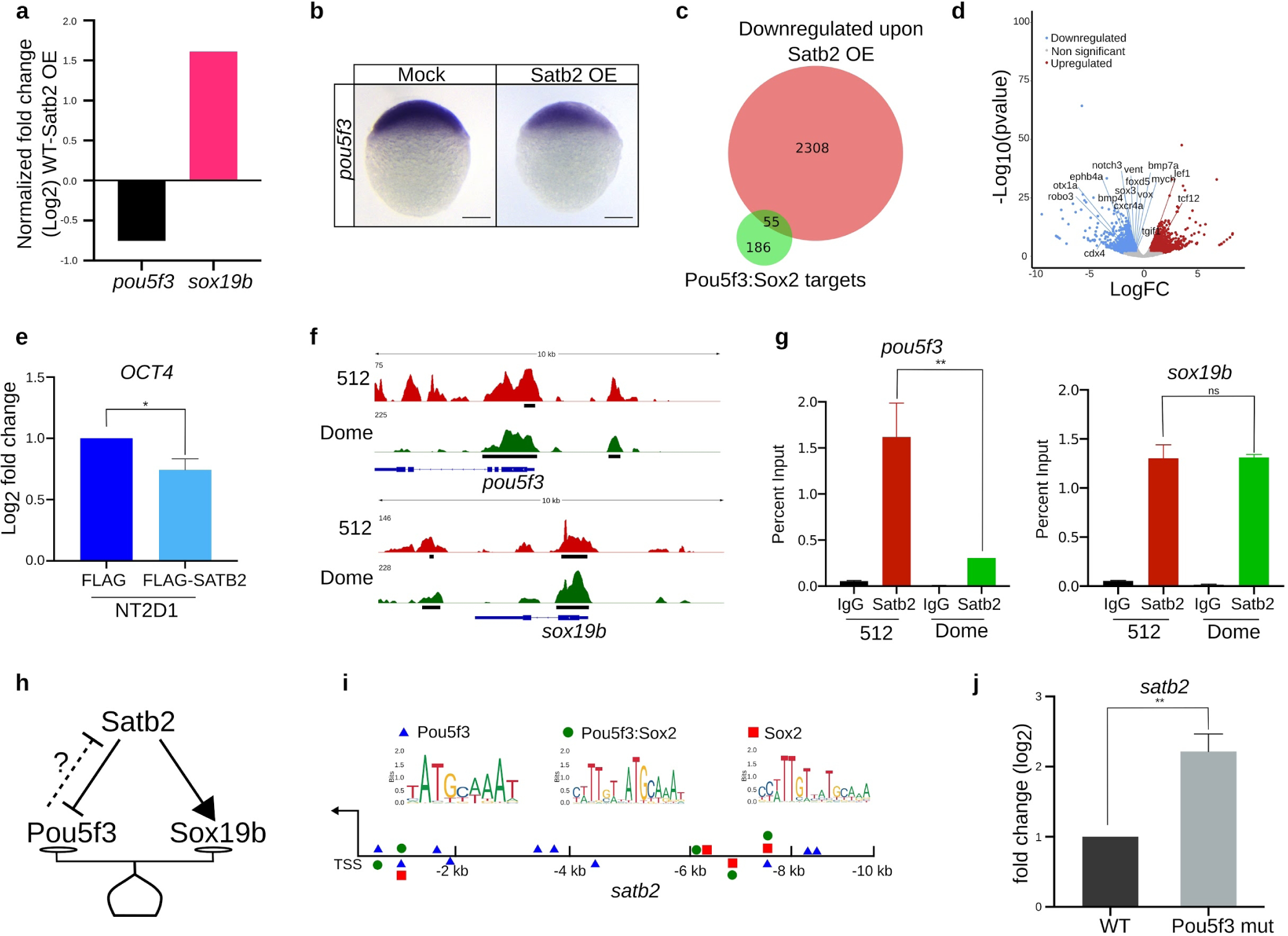
Maternal Satb2 functions during ZGA by regulating pioneer factors. **a**, Relative gene expression for *pou5f3* and *sox19b* as determined by RNA-seq analysis (FDR < 0.05). **b**, Lateral view of zebrafish embryo at 4 hpf for whole-mount in situ hybridization using RNA probe against *pou5f3* in control and Satb2 overexpressing embryos (n=24). **c**, Venn diagram analysis showing the overlap between negative targets of Satb2 and known genomic targets of Pou5f3 and Sox2. **d**, Volcano plot of differentially expressed genes upon overexpression of Satb2 analyzed at 4.5 hpf. DE genes with FDR < 0.1 and log2 Fold change > +/− 0.58 are coloured as red for upregulated, blue for downregulated and grey for unchanged or non-significant genes. Key Pou5f3: Sox2 target genes are labelled. **e**, Quantitative RT-PCR analysis of *OCT4* (human homologue of *pou5f3*) expression in NT2D1 cells upon overexpression of SATB2 as compared to FLAG overexpression. * signifies P-value < 0.05 as calculated by the Student’s two-tailed t-test, N=3. **f**, IGV snapshot of Satb2 ChIP-seq at 512 cells (red) and Dome stage (green) on genomic loci of *pou5f3 and sox19b*. **g**, Quantitative relative enrichment (ChIP-qPCR) represented as percent input using isotype-matched IgG and anti-Satb2 antibody for genomic locus (TSS) of *pou5f3* and *sox19b*. ** indicates P-value < 0.001 as calculated by the Student’s two-tailed t-test, N=2. **h**, The predictive regulatory model between maternal Satb2 and pioneer factors during the establishment of ZGA. **i**, Schematic for *satb2* promoter displaying motif sites marked for Pouf53, Pou5f3: Sox2 and Sox2. **j**, qPCR analysis for *Satb2* expression at 4.5 hpf in *pou5f3* mutant embryos. ** signifies P-value < 0.01 as calculated by the Student’s two-tailed t-test, N=3.

To verify if the negative regulation is mediated by direct binding of Satb2, we performed ChIPseq at the pre-MBT stage (512 cells) and post MBT stage (dome) (Supplementary Fig. 7b). We observed strong enrichment of Satb2 at the TSS of both *pou5f3* and *sox19b* (Fig. 6f). However, ChIP-qPCR analysis showed reduced Satb2 occupancy on the genomic *pou5f3* locus from pre- to post-MBT but not on the promoter of *sox19b* (Fig. 6g). Such dynamic occupancy pattern was also observed for other negative targets of Satb2 including *ta*, *bmp4* and *gsc* (Supplementary Fig. S7c). Altogether, these results support that depletion of maternal *satb2* is essential for sustained expression of the pioneer factor *pou5f3* and in turn the onset of ZGA. However, we were curious as to how the maternal pool of *satb2* is regulated at the onset of ZGA. Since the Pou5f3, Nanog and SoxB1 act as pioneer factors, we asked if there is a feedback regulation between these factors and Satb2 (Fig. 6h). To test this, we first scanned the *satb2* promoter (up to −10 Kb) for binding motifs of Pou5f3:Sox2 complex, Pou5f3 alone and Sox2 alone. A number of potential binding sites were observed for these pioneer factors within a 2 Kb window upstream to TSS (Fig. 6i). Previously published ChIPseq data at post-MBT stages supported this observation (Figure S7E). To establish the functional relevance of the binding of Pou5f3, we performed expression analysis of *satb2* using qRT-PCR in *pou5f3* mutants. Significant upregulation of *satb2* thus supports the presence of possible negative feedback regulation between Satb2 and the pioneer factor Pou5f3 (Fig. 6j).

### Early developmental regimes established using human embryonic stem-like cells recapitulate biphasic nature of *SATB2* expression

Temporal analysis of Satb2 expression revealed an interesting biphasic pattern that translated into bimodal regulation. We wondered if this characteristic pattern can be mimicked using pluripotent NT2D1 embryonic cells. Upon retinoic acid stimulation NT2D1 cells differentiate into neuronal progenitors^45^. We found that while *SATB2* is readily detectable in undifferentiated cells, *SATB2* levels decrease during transition stages (day 3 post RA treatment). Interestingly, we also observed an increase in *SATB2* expression as cells differentiated into neuronal progenitors, characterized by an increase in *PAX6* expression (Supplementary Fig. 8a). Next, we analyzed the transcriptome of H9 human ES cells differentiation series^46^. Even in this scenario, *SATB2* is initially transcribed in the pluripotent cells, and gets down regulated upon initiation of differentiation between day 2 to day 6. *SATB2* transcription is restored during later stages (day 8 and beyond) when neuronal fate specification ensues (Supplementary Fig. 8b).

### Stage-specific occupancy analysis of Satb2 reveals the underlying mechanisms for the differential activity of maternal and zygotic Satb2

Our results highlight the contrasting nature of maternal versus zygotic Satb2 function during early embryogenesis. Maternal Satb2 primarily functions as a repressor while zygotic Satb2 activates neurogenesis programs. Such disparate behavior could be attributed to two potential mechanisms: (1) Differential occupancy at the target loci and/or (2) Stage specific distinct interacting protein partners. Towards this, we evaluated Satb2 occupancy by revisiting our ChIPseq datasets from 512 cells stage to 14 ss. Interestingly, we observed a substantial shift in the ratio of non-promoter to promoter-bound regions by Satb2 (Fig. 7a). Maternal Satb2 occupies more of the non-promoter regions (67% intergenic and 10% promoter) at the pre-MBT stage, whereas zygotic Satb2 localizes substantially to the promoter regions (54% intergenic and 36% promoter).

**Fig. 7:**
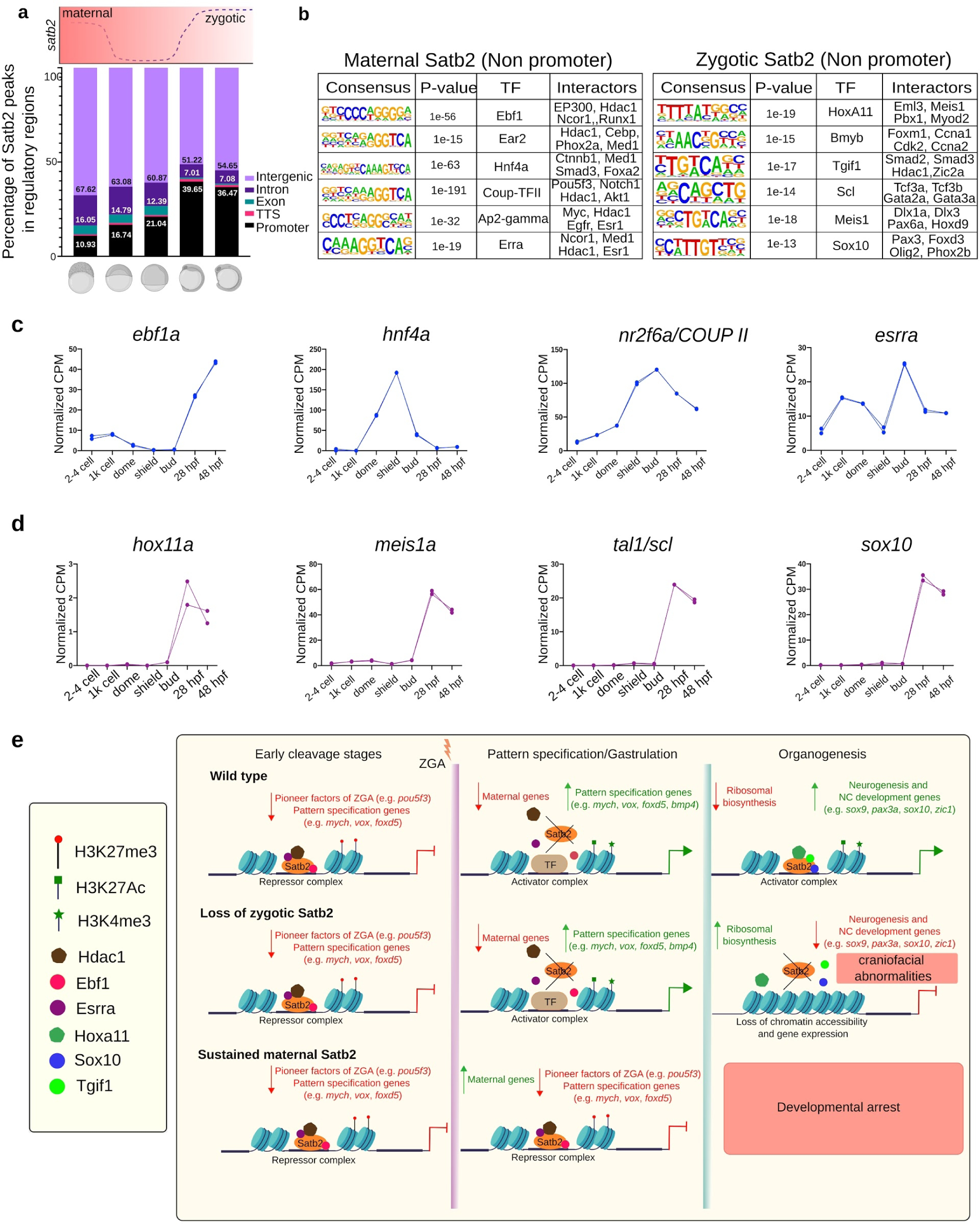
Differential mechanisms employed by maternal and zygotic Satb2 to perform contrasting functions during embryogenesis. **a**, Genomic distribution of Satb2 bound regions across developmental stages highlighting a shift in the ratio of non-promoter to promoter bound peaks. Schematic above the bar plot represents the expression pattern of *satb2* throughout early embryogenesis. **b**, TFBS for transcription factors at maternal (512 cell stage) and zygotic (14 ss) Satb2. Known Interactors for each transcription factor are enlisted. **c**, Line graph depicting mRNA expression dynamics for putative interactors of maternal Satb2; *ebf1a*, *hnf4a*, *nr2f6a* and *esrra* across developmental stages as indicated on the x-axis. **d**, Line graph depicting mRNA expression dynamics for putative interactors of zygotic Satb2 namely, *hox11a*, *meis1a*, *tal1*, and *sox10,* across developmental stages as indicated on the x-axis. **e**, Schematic summary highlighting differential functions of Satb2 during various stages of zebrafish embryogenesis.

Next, we performed consensus binding analysis for non-promoter and promoter-bound regions at 512 cell stage and 14 ss to identify putative interactors of Satb2. Maternal Satb2 occupied regions were enriched for binding sites of Ebf1, Ear2 (nr2f6), Hnf4a, Coup-TFII (Nr2f2), Ap2-gamma (Tfap2c) and Erra. Among these, Ebf1, Ear2, Coup-TFII proteins have been widely characterized for repressive activity during B cell and NC differentiation^47, 48^ (Fig. 7b). In contrast, zygotic Satb2 binding regions show enrichment for Hoxa11, Bmyb, Tgif1, Scl, Sox10 and Meis1 all essential factors for NC differentiation and neurogenesis^28,34,39,49,50,51^.

Expression dynamics of these transcription factors highlights putative stage-dependent association with Satb2 (Fig. 7c,d). Using the STRING database and published literature, we extracted putative interactors of these proteins. Potential interactors of maternal Satb2 display propensity to be associated with co-repressors such as Hdac1, Myc, and Med1^52–54^ whereas interactors of zygotic Satb2 are likely to partner with the positive regulators of neurogenesis including Pitx1, Tcf3a, Dlx3a, Tgif1, Olig2 and Zic2a^34, 55–59^. This analysis argues that contrasting stage-dependent function of Satb2 is a consequence of qualitatively distinct, unique sets of binding partners (Fig. 7e). However, future experiments will be necessary to test this idea.

## Discussion

Early embryonic development is a culmination of carefully calibrated gene expression patterns orchestrated in a spatiotemporal manner. Typically, this outcome is achieved by establishing tissue specific GRNs. How the GRNs are initiated, maintained and executed has been a major focus of inquiry among developmental biologists. Consequently, we have learnt a great deal about regulation of tissue specific gene expression leading to cell fate specification. In recent years the focus has gradually shifted to elucidation of genome-wide molecular changes that engineer critical cellular/developmental landmarks such as ZGA or early determination of cell type identity. However, how the global changes in genome architecture are mechanistically connected to individual loci has still remained mysterious. Equally unknown are the individual players that participate as crucial molecular links between these processes. Here, we have investigated the function of the chromatin organizer protein Satb2 in developmental transitions during early vertebrate embryogenesis.

Studying Satb2 in this regard is especially pertinent as it appears to be the most ancient member of its cohort across evolutionary scale and its function is required during embryogenesis for proper specification of NCCs, a cell type critical for craniofacial development. Indeed, the zebrafish *satb2* mutants generated in this study recapitulated phenotypes seen in corresponding aberrations in mice and also resembles the human genetic conditions induced by the disruption of *SATB2* locus. Emboldened by this similarity we embarked on a detailed analysis of *satb2* mutant embryos using functional genomics. We simultaneously employed analogous samples using mouse tissues and human cell lines and, thus far, several observations of broad significance have emerged.

First, Satb2 binds to multiple cis-regulatory sequences in a genome-wide manner. Moreover, Satb2 can act both as a repressor and as an activator of transcription. Consistently, we have identified several positive and negative targets of Satb2 through transcriptome, genome-wide occupancy and chromatin accessibility analysis. For instance, Satb2 activates transcription of the effectors of NC specification while it downregulates components of ribosomal biosynthesis. In future, it would be of interest to investigate the functional relationships between groups of such gene products. In fact, such cross regulatory interactions that fine tune gene expression via feed-forward and feed-back mechanisms is an established hallmark of embryonic development.

The most remarkable finding that emerged from this study is the functional dichotomy between the maternal and zygotic Satb2. By analyzing the maternal pool of Satb2 we were able to trace back it’s activity to the very early stages of embryonic development. Maternally deposited Satb2 diminishes at the onset of ZGA before resuming function during zygotic development including organogenesis. Intrigued by this biphasic nature of *satb2* expression, we analyzed the function of maternal Satb2 by employing morpholino based knockdown and overexpression. Surprisingly, analysis of the pluripotent embryonic cells uncovered that maternal Satb2 acts as a transcriptional repressor of zygotic genes involved in patterning. It is also noteworthy that under these circumstances metabolic pathways appeared to be upregulated which could be either a direct or an indirect consequence of perturbation in ZGA. This is in sharp contrast to the function of zygotic Satb2 suggesting that the biphasic expression of Satb2 is instrumental in manifesting bimodal, contrasting molecular functions. Perturbation of ZGA by Satb2 overexpression resulted in severe developmental defects. Consistent with our observations at 14 ss, we found maternal Satb2 also regulates local chromatin landscape of target genes by modulating histone modification occupancy profiles. Reprogramming of histone modification patterns is essential for successful onset of ZGA^60, 61^. Together these studies indicate that maternally deposited H3K27me3 is depleted prior to onset of ZGA with concomitant increase in H3K4me3 and H3K27Ac. Interestingly, we did not observe significant perturbation in chromatin accessibility. Thus, we propose that during cleavage stages, Satb2 brings about transient chromatin modulations rather than generating lasting effects through active chromatin remodeling.

The bimodal activity of Satb2 poses two major questions. What regulates expression of Satb2 during embryonic development? And how does Satb2 acquire differential regulatory potential? BMP and SHH signaling regulate the expression of *satb2*^10, 20^. Since these signalling pathways are dormant during early cleavage stages and thus the upstream regulators of maternal Satb2 needs further investigation. We have discovered a novel negative feedback regulation between Satb2 and pioneer pluripotency factors such as Pou5f3. Notably, we observed that Satb2 regulates Pou5f3 and Sox19b in contrasting ways to maintain gene expression homeostasis necessary for ZGA. Such a mechanistic interplay has been proposed by Gao *et al*., providing novel insights into how the balance between Pou5f3 and Sox19b is essential for efficient ZGA^62^. Pou5f3 has been implicated in regulating ZGA in zebrafish and humans but not in mice^3, 4, 63^. We propose that Satb2 potentially act as a determinant of organism specific GRNs by regulating one or more of the “Yamanaka” factors during the process of ZGA.

Next, we sought to analyze underlying mechanisms for contrasting functions of maternal and zygotic Satb2. Stage-dependent ChIP-seq analysis revealed a dynamic shift from the non-promoter- to promoter-bound fraction of Satb2. Based on the motif analysis we propose that Satb2 indeed has the potential to form a complex with multiple partners and coordinate gene expression based on the expression dynamics of the interacting partners during embryogenesis (Fig. 7c,d). Such mechanisms have been proposed for bimodal activity of NC fate determinant Foxd3^64^. Moreover, the gene regulator function of SATB1 is known to be governed in bimodal fashion via interaction partners in T cells^65^.

In summary, we have characterized temporally regulated context-dependent functions of Satb2 (Fig. 7e). Our studies have unraveled involvement of Satb2 as a gatekeeper during major regulatory events throughout early vertebrate embryogenesis. This study also underscores Satb2 as a bonafide member of a group of proteins that wear multiple hats and function in a similar biphasic and bimodal manner during body plan determination.

## Author Contributions

S.J.P. conceived the study, designed experiments, interpreted data, generated Crispr mutants, performed all zebrafish embryology experiments with assistance from other authors, performed RNA-seq, ChIP-seq, ATAC-seq, scRNA-seq and associated data analysis, generated figures, and wrote the manuscript. P.C.R. performed phylogenetic analysis, protein domain architecture analysis, and performed bulk RNA-seq analysis. M.Sm. designed and assisted in overexpression experiments. A.S. performed micro CT scans, assisted in scRNA-seq and ATAC-seq. K.S. cloned 3xFLAG-Satb2 and assisted in experiments. M.S.O. performed alcian blue staining and WISH. R.S. performed SATB2 overexpression experiments in NT2D1. M.P. Assisted in stage-dependent ChIP-seq experiments. O.D. performed WISH and morpholino mediated knockdown. Y.T. generated plots for Satb2 OE mRNA-seq analysis. R.K.M. contributed resources. G.D. edited the manuscript and provided expertise in evolutionary perspectives. A.J.G designed GLT experiments and provided expertise in *pou5f3* mutants and ZGA regulation. M.S. provided expertise in zebrafish experiments, design of morpholino experiments and provided resources. C.-P.H. designed experiments, interpreted data, provided resources, edited manuscript, obtained funding, and supervised the study. S.G. conceived the project, designed experiments, interpreted data, wrote manuscript, obtained funding and other resources, and supervised the entire study. All authors read and approved the final manuscript.

## Acknowledgments

We are grateful to members of C.-P.H. and SG lab for discussions. Authors thank Shubha Tole for providing embryonic mouse tissues. Authors would like to thank Satyajeet Khare, Vanessa Barone, Jyothish S., Greg D’silva, Shalini Mishra, Yoshita Bhide, and Keshav Jha for assistance in experiments. We would also like to thank Chaitanya Dingare for valuable suggestions. We thank Diana Pinhiero and Alexandra Schauer for critical reading of early versions of manuscript. This work was supported by the Centre of Excellence in Epigenetics program of the Department of Biotechnology, Government of India Phase I (BT/01/COE/09/07) to S.G. and R.K.M., and Phase II (BT/COE/34/SP17426/2016) to S.G., and JC Bose Fellowship (JCB/2019/000013) from Science and Engineering Research Board, Government of India to S.G., DST-BMWF Indo-Austrian bilateral program grant to S.G. and C.-P.H. S.J.P. was supported by Fellowship from the Council of Scientific and Industrial Research, India and travel fellowship from the Company of Biologists, UK. P.C.R. is supported by the Early Career Fellowship of the Wellcome Trust-DBT India Alliance (IA/E/16/1/503057). A.S and R.S. were supported by CSIR India. M.S. was supported by core funding from the Tata Institute of Fundamental Research (TIFR 12P-121).

## Competing interests

The authors declare no competing interests.

## Methods

### Key Resource table

| Reagents/Resources | Source | Identifier |
| --- | --- | --- |
| **Antibodies** |  |  |
| Anti-H3K27Ac | Abcam | cat# ab4729 |
| Anti-H3K27me3 | Millipore | cat# 07-449 |
| Anti-H3K4me3 | Abcam | cat# ab8580 |
| Anti-gamma-Tubulin | Sigma | cat# T6557 |
| Anti-FLAG-M2 | Sigma | cat# F3165 |
| Anti-SATB2 | Abcam | cat# ab34735 |
| Anti-zebrafish Satb2 | This study | cat# NA |
| Anti-Human Satb2 | This study | cat# NA |
| Anti-DIG-AP Fab | Roche | cat# 1093274 |
| Anti-Rabbit IgG | Invitrogen | cat# 31235 |
| Anti-Rabbit-HRP conjugate | Bio Rad | cat# STAR124P |
| **Critical Commercial Reagents** |  |  |
| Quantseq 3' mRNA seq library kit | Lexogen | cat# 015.96 |
| SENSE mRNA-Seq Library Prep Kit | Lexogen | cat# 001.96 |
| NEB ultra II DNA library kit | NEB | cat# E7645L |
| Nugen ultra II ovation DNA kit | Nugen | cat# 0344NB-A01 |
| Single Cell 3' Reagent Kits V3 | 10X Genomics | cat# PN-1000092 |
| Hiprep PCR cleanup kit | Magbio | cat# AC-60050 |
| SPRIselect | Beckman<br>Coulter | cat# B23318 |
| NextSeq 500/550 High Output Kit | Illumina | cat# 20024906 |
| v2.5 (75 cycles) |  |  |
| NextSeq 500/550 Mid Output Kit<br>v2.5 (150 cycles) | Illumina | cat# 20024904 |
| Dynabeads Protein A | Invitrogen | cat # 10002D |
| Dynabeads Protein G | Invitrogen | cat# 10004D |
| TDE1 Tn5 | Illumina | cat# 15027865 |
| TD Buffer | Illumina | cat# 15027866 |
| **Experimental Models:**<br>**Organisms/Strains** |  |  |
| Zebrafish: Tuebingen wild-type | MPI-CBG<br>Dresden | NA |
| Zebrafish: *satb2*-STOP | This study | *satb2*-E1-STOP |
| Zebrafish: MZ*pou5f3* | Burgess et al.<br>2002 | *spg* |
| Mouse: SWR/J | The Jackson<br>Laboratory | Stock# 000689 |
| **Recombinant DNA clones** |  |  |
| pCS2-zf.3xFLAG-*satb2* | This study | NA |
| pCS2-zf.*satb2* | This study | NA |
| pCS2-zf.mismatch-*satb2* | This study | NA |
| pET28b-6XHis-zf.*satb2*-STREP | This study | NA |
| CMV9-3XFLAG-Hu.*SATB2* | This study | NA |
| pET28b-Cas9-His | Gagnon et al.<br>2014 | Addgene# 47327 |
| **Deposited Data** |  |  |
| mRNA-seq | This study | NCBI BioProject ID PRJNA657343 |
| Single cell RNAseq | This study | NCBI BioProject ID PRJNA657343 |
| ChIP-seq | This study | NCBI BioProject ID PRJNA657343 |
| ATAC-seq | This study | NCBI BioProject ID PRJNA657343 |
| **Software and Algorithms** |  |  |
| Trimmomatic 0.39 | Bolger et al.<br>2014 | https://github.com/timflutre/trimmomatic |
| BWA | Li H. and<br>Durbin R.<br>(2009) | http://bio-bwa.sourceforge.net/ |
| STAR2.4.2 | Dobin et al.<br>2013 | https://github.com/alexdobin/STAR |
| BBmap | Bushnell B.<br>2014 | https://sourceforge.net/projects/bbmap/ |
| DeepTools | Ramirez et al.<br>2016 | https://github.com/deeptools/deepTools |
| HOMER (v.4.4) | Heinz et al.<br>2010 | http://homer.ucsd.edu/homer/index.html |
| MACS2 (v2.1.0) | Zhang et al.<br>2008 | https://github.com/taoliu/MACS |
| FeatureCounts (v1.4.6-p4) | Liao et al. 2014 | http://bioinf.wehi.edu.au/featureCounts |
| edgeR | Robinson et al.<br>2010 | https://bioconductor.org/packages/release/bioc/html/edgeR.html |
| Diffbind | Stark and<br>Brown 2011 | https://bioconductor.org/packages/release/bioc/html/DiffBind.html |
| Seurat 3.0 | Butler et al.<br>2018 | https://satijalab.org/seurat/ |
| NucleoATAC | Schep et al.<br>2015 | https://github.com/GreenleafLab/NucleoATAC |
| Fiji | Schindelin et al.<br>2012 | https://fiji.sc/ |
| GraphPad Prism | GraphPad<br>software | https://www.graphpad.com/scientific-software/prism/ |
| R version 3.6.1 | The R Project<br>for Statistical<br>Computing | https://www.r-project.org/ |
| BioRender | BiorRender | https://biorender.com/ |

### Experimental models

Zebrafish (Danio *rerio*) were maintained as described previously^66^. All the experimental procedures were carried out in accordance with the guidelines from the institutional animal ethics committee at IISER Pune and IST Austria. Embryos were raised at 23-31°C in E3 medium and staged as described previously^67^. Experiments with mice models (*Mus musculus*) were carried out at TIFR, Mumbai adhering to Institutional ethics guidelines.

### Phylogenetic analysis of SATB proteins

Homologous sequences for SATB proteins were extracted from GeneTree data from Ensembl (http://www.ensembl.org/Multi/GeneTree/Image?gt=ENSGT00390000008096) and were further curated using NCBI BLASTP in selected organisms for their true copy in their genome to avoid inclusion of splice variants. Curated sequences were aligned using MUSCLE using default parameters and alignments were trimmed using trimAl on auto mode which are embedded in the online server (https://ngphylogeny.fr)^68^. A phylogenetic tree was generated using randomized accelerated maximum likelihood method (RAxML) method available at CIPRE Science Gateway server^69^. Here, PROTCAT was used as a substitution model with DAYHOFF substitution matrix. DVE CAEL was used as an outgroup and 1000 bootstraps were performed for the analysis. The resulting phylogenetic tree was visualized using iTOL and tree branches are colour coded bases on different groups^70^. The representative animal’s silhouette images were collected from (http://phylopic.org/) and used in annotation.

### CRISPR/Cas9 mutant generation

Synthetic oligonucleotides targeting exon1 of *satb1* gene using CRISPR-Cas9 method were designed using CHOPCHOP and cross-validated using CRISPRscan^71, 72^. Oligonucleotides required to generate sgDNA synthesized commercially as listed below (IDT, USA).

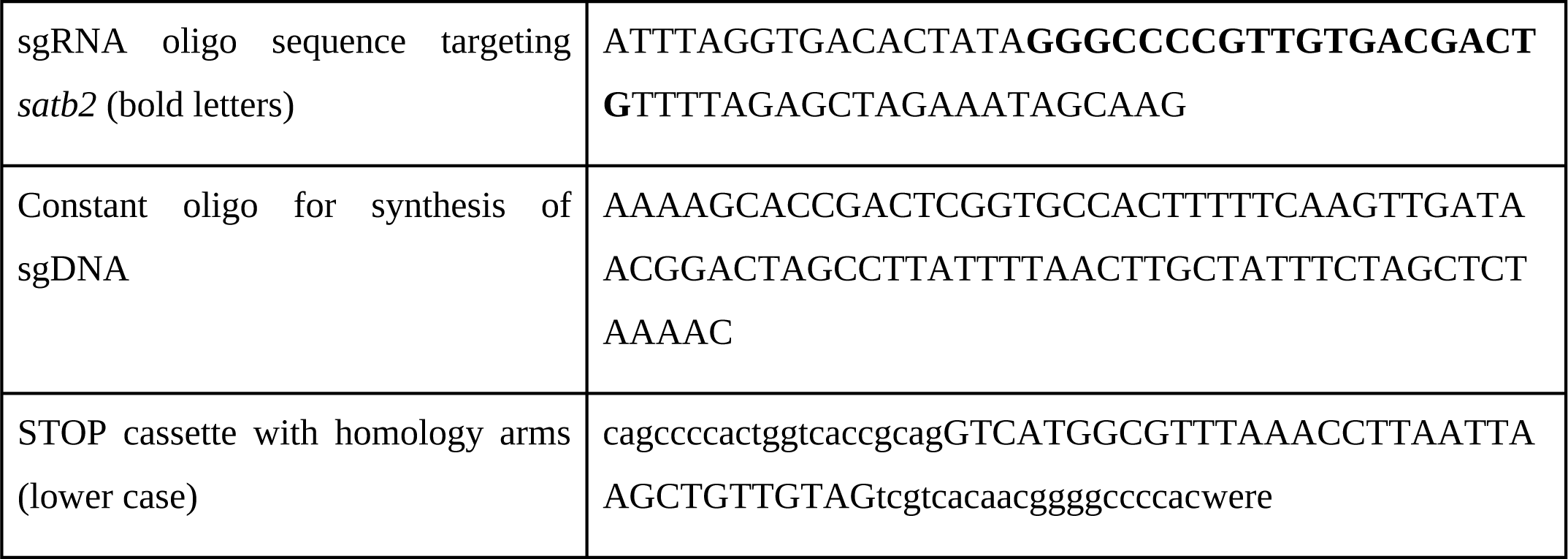

sgDNA template was generated using gene-specific oligonucleotides and constant oligonucleotide by polymerase chain reaction for 30 cycles using Q5 high fidelity polymerase (NEB, USA). PCR product was further purified using Magbio Hiprep PCR cleanup system (Magbiogenomics, USA) and quantified using Nanodrop 2000. sgRNA was synthesized *in vitro* as per the suggested protocol using Ampliscribe T7 Flash Transcription kit (Lucigen, USA). 6xHis-Cas9 protein was synthesized as described previously^25^. Briefly, plasmid coding for 6x-His-Cas9 were transformed in Rosetta DE3 Novagen competent cells (Merck Millipore, USA). Protein production was induced using autoinduction medium (Formedium, UK) for 36 hrs at 18°C with constant shaking at 200 RPM. Bacterial cells were harvested and lysed in 20 mM Tris pH 8.0, 30 mM Imidazole, 500 mM NaCl followed by sonication with 20 secs ON 20 secs OFF pulses for 30 mins using a probe-based sonicator, Sonics Vibra cell (Sonics, USA). Lysates were cleared using high-speed centrifugation and incubated with HisPur cobalt resin (Thermo scientific, USA) for 1 hr. Beads were further washed three times with lysis buffer and eluted in 20 mM Tris pH 8, 200 mM Imidazole, 500 mM NaCl. Eluted fractions were further dialyzed in 20 mM Tris, 200 mM KCl, 10 mM MgCl_2_ and stored as 5 ul aliquots at −80 for further use.

The sgRNA-Cas9 protein complex (sgRNA 200ng/ul, Cas9 600ng/ul and STOP cassette 1uM) was injected at the one cell stage of zebrafish embryos. Founders were obtained by genotyping using oligonucleotides flanking the target site and maintained as heterozygous.

### Genotyping of Satb2 mutants

The embryos used for all the experiments were a result of satb2^+/−^ incrosses. As a result, genotyping of the embryos after every experiment was necessary. DNA from embryos/larvae at desired stages of development were extracted in 50ul of genomic DNA extraction buffer containing 10ug proteinase K at 55°C for 16 hrs (10 mM Tris-HCl pH 8.0, 50 mM KCl, 0.3% Tween-20 and 0.3% NP40). Proteinase K was inactivated by incubating at 95°C for 20 mins followed by snap chilling. For adult fish, genotyping was performed using fin clips as described^66^. PCR was carried out using forward primer: 5’GGAGGAGAGAGTCCTCGACTG3’, Reverse primer: 5’GTTGCAGCATGTTTCAGATGAT3’ with paq polymerase 5000 (Agilent, USA) and resulting PCR products were electrophoresed on 3% agarose gel. Gel images were captured using G:Box gel documentation system (Syngene, USA).

### Micro-CT image acquisition and analysis

*Satb2^+/−^* fish were incrossed and were grown until 4 months post fertilization. Fish were fin clipped and genotyped using the protocol described above. Screened fish (N=5 per group) were fixed in a 3.7% paraformaldehyde and 1% glutaraldehyde in 1X PBS solution, fish were fixed for 72 hours at room temperature and then transferred to 1% of fixative solution for storage. Fish were imaged first using SLR cameras and then were imaged using Quantum GX microCT imaging system (Perkin Elmer, USA). Image acquisition settings were as follows, Temperature: 23°C, Humidity: 57%, Voltage: 90 kVp, Current: 88 uA, X-Ray dosage: 449 mGy, FoV: 36mm, Recon: 36 mm, Voxel size: 72 um, Scan mode: high resolution, Time: 4 minutes. Images were further processed with ImageJ (FIJI) for adjusting brightness and contrast.

### Alcian blue - Alizarin red double staining

Alcian blue: Alizarin red acid-free double staining was performed as described previously^73^. All solutions were stored and all steps were carried out at room temperature. Briefly, embryos from satb2^+/−^ incrosses were grown to 15 dpf and fixed with 4% PFA for 2 hours. After dehydration for 10 min in 50% ethanol, larvae were stained overnight in a double staining solution made from a mixture of Alcian blue: Alizarin red stock solution in a 100:1 ratio respectively. Alcian blue stock solution for cartilage staining contained 0.1% Alcian blue powder (Sigma Aldrich) in 70% ethanol and 60mM MgCl2 final concentrations. Alizarin red (Sigma Aldrich) stock solution for bone staining was made at a final concentration of 0.5% in MQ water. After staining, the solution was removed by a quick rinse with MQ water followed by washes with 1:1 solution of 3% H_2_O_2_ and 2% KOH for 20 min with the tube caps kept open. Larvae were then cleared with successive washes with increasing concentration of glycerol (20-50%) and 0.25% KOH solutions. Larvae were finally stored in a 50% glycerol and 0.1% KOH storage solution until imaging. At the time of imaging, larvae were indexed and temporarily mounted with the help of 75% glycerol and 0.1% KOH solution and imaged on an Olympus light microscope. After 3 subsequent washes with PBST to remove the bone stain, selected larvae were genotyped for representation.

### 3’mRNA gene expression assay and differential gene expression analysis

Single embryos were harvested in 100ul RNA iso-plus total RNA extraction reagent (DSS Takara, India) at 80% epiboly, 6 somites and 14 somites stage of zebrafish embryonic development. Samples were homogenized by vigorous vortexing to assure complete lysis followed by addition of 15 ul of CHCl3. Aqueous layer was separated by centrifugation at 12000 g for 15 mins at 4°C. The aqueous layer was collected in a separate tube and an equal volume of isopropanol containing GlycoBlue coprecipitant (Invitrogen, USA) was added. The aqueous fraction was stored at −20 until further processing. Meanwhile, the organic fraction was processed for DNA extraction by precipitating with 150ul of 100% ethanol and incubated for 10 mins at RT. Samples were centrifuged for 10 mins at high speed and the pellet was washed with 70% ethanol (twice). After drying, DNA pellet was resuspended in Low TE and used for genotyping as mentioned before. Once genotype for the single embryo is confirmed, corresponding RNA fractions were processed further. We pooled 2-3 samples to get a sufficient amount of RNA to prepare libraries.

Extracted RNA samples were quantified using Nanodrop 2000 and RNA integrity was determined using Agilent bioanalyzer 2100. Only samples with RIN > 8 were used to prepare sequencing libraries using Quantseq 3’ mRNA seq library kit (Lexogen GMBH, Austria) as per the instructions from manufacture. Briefly, 300 ng of RNA was used for polyA capturing and subjected to first and second-strand synthesis. All cDNA samples were amplified for 10-15 cycles depending on cycle number estimation by qPCR (PCR add on kit, Lexogen GMBH, Austria). Amplified libraries were purified using two rounds of 0.8x volume of Hi-prep PCR purification kit. The concentration of libraries was estimated using the Qubit DNA HS system. Finally, all the libraries were pooled and subjected to high throughput sequencing using 76 bp SE chemistry on Nextseq 550.

Sequencing reads were trimmed for quality using Trimmomatic and aligned to daRer10 genome assembly using STAR aligner. Counts for each gene feature was estimated using the FeatureCounts package from Rsubread. Differential expression analysis was performed for replicates using EdgeR. Volcano plot for significantly differentially expressed genes was generated using the PlotVolcano tool from Galaxy server. Gene ontology analysis for upregulated and downregulated genes were performed using a web-based tool, gProfiler^74^.

### Synthesis of riboprobes for Whole-mount in situ hybridization

Synthetic oligonucleotides against target mRNAs (listed below) with T7 RNA polymerase promoter sequence on the 5’ end of the reverse primers were obtained commercially (Sigma Aldrich, India). cDNA templates for corresponding stages were used for PCR amplification. PCR reactions were purified using NEB Monarch DNA Gel Extraction Kit or HighPrep PCR Clean-up System beads. Further, Riboprobes were synthesized using Roche DIG RNA labelling kit at 37°C for 3 hrs followed by DNAse treatment for 15 mins. Probes were further purified using Bio-Spin 30 Tris Columns (Biorad, Germany). The digoxigenin-labelled probes were 400-700 nucleotides in length and stored at −20 deg C at 1 ug/ml.

For *chrd* WISH experiments amplified region was cloned into TOPO II dual promoter vector (Thermo scientific, USA). To generate antisense probe, plasmid was linearised using BamHI and probe synthesis reaction was carried out using T7 RNA polymerase promoter as described above.

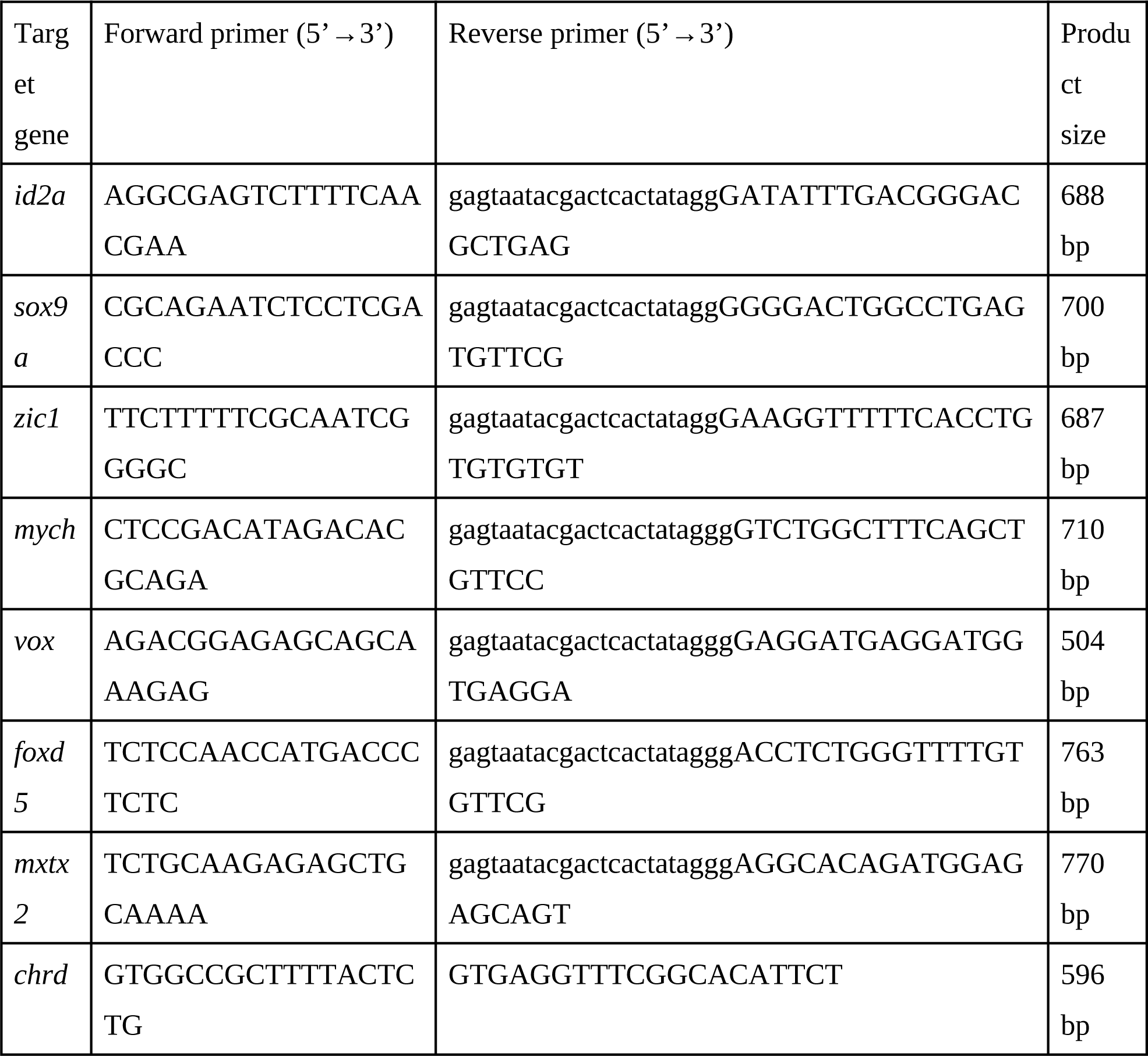
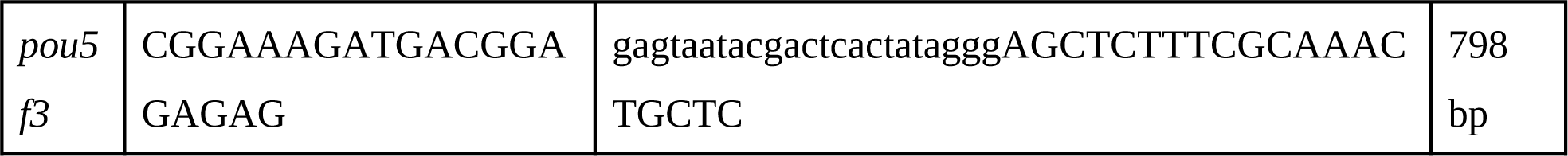

### Whole-mount In situ hybridization analysis

Whole-mount In situ hybridization (WISH) was performed as described previously^75^. Fertilized embryos from satb2^+/−^ incross were raised until the desired stage of development was reached. Embryos were fixed in 4% paraformaldehyde in 1x PBS overnight at 4°C with gentle rocking. Next day, embryos were hand dechorionated with the pair of fine forceps and incubated in 100% MeOH overnight at −20°C. Embryos were further downgraded in methanol: PBST solution (0.1% Tween-20 in 1x PBS pH 7.4). Following successive rehydration to PBST, embryos were treated with 5ug/ul proteinase K solution in PBST for 2 min for 14ss stage embryos. The reaction was stopped by a subsequent 20 min incubation in 4% PFA. For Dome stage, embryos were not treated with proteinase K. After washes in PBST, samples were pre hybridized for 2-5 hours and incubated overnight at 70°C in approximately 500 ng of the respective probe. Next day, embryos were washed with SSC buffers and further blocked in blocking buffer (2% v/v normal goat serum, 2mg/ml BSA in PBST) for 3-4 hours at room temperature followed by overnight incubation at 4°C in anti-DIG-AP Fab fragments (1:5000) (Roche 1093274). Embryos were washed with PBST followed by an alkaline tris buffer. Staining was performed with BM Purple substrate (Roche). After staining, the color reaction was stopped using a stop solution containing 20% Tween-20 and 05M EDTA in 1x PBS (pH 5.5). Embryos were further washed with PBST and stored at 4°C until imaging. Embryos for imaging were mounted on agarose moulds and imaged using Z-stacking mode on a Leica DFC450C microscope. Post imaging, embryos were genotyped using the standard protocol discussed above. Representative images were further processed using ImageJ for adjusting brightness and contrast.

### Antibody production

All the procedures were performed as per the approved guidelines from the ethical committee at national toxicology centre (NTC), Pune. To generate polyclonal antibodies, anti-zebrafish Satb2 (c-term): CIPSSGAEENPQANTGSGNNGP and anti-human SATB2 (c-term): CQQSQPAKESSPPREEAP peptides were synthesized commercially (Apeptide, China). Antibodies were produced in New Zealand white Rabbits, as per the protocols from the laboratory of Tony Hyman, MPI-CBG with modification as below (https://hymanlab.mpi-cbg.de/hyman_lab/general/). Briefly, the required amount of peptides were conjugated with KLH using glutaraldehyde followed by subsequent dialysis to remove glutaraldehyde. Conjugated peptides were mixed with Freud’s complete adjuvant (Sigma Aldrich) for the first immunization. Rabbits were immunized intradermally. Further, after every 21 days, rabbits were immunized using peptides mixed with Freud’s incomplete adjuvant until sufficient titre for the antibody was obtained.

Antisera was purified by a peptide affinity column prepared using sulfoLink coupling resin according to the manufacturer’s instructions (Thermo) and stored in 50% glycerol solution at −20°C.

### Molecular cloning of expression plasmids

For zebrafish Satb2 over-expression studies, full length *satb2* was amplified from cDNA samples generated using High capacity cDNA synthesis kit (Thermo scientific, USA) from 48 hpf wild type TU embryos. Resulting PCR product was purified using Monarch PCR purification kit (NEB, USA) and were cloned into pCS2+ vector using restriction enzymes BamH1 and SnaB1 based cloning to generate pCS2-Satb2. To generate 3xFLAG-Tagged construct, oligo containing 3xFLAG sequence at the 5’ end of the forward primer was used for PCR using pCS2-Satb2 plasmid as a template and clone was obtained using gibson assembly. To generate morpholino resistant clones, oligo containing 7 mismatches spanning the first 24 bases of the coding region were used for amplification using pCS2-Satb2 plasmid as a template.

For recombinant expression of zebrafish Satb2 protein, coding sequences were amplified using Q5 high fidelity DNA polymerase (NEB, USA) and cloned into 6xHis-PET28-Strep vector (Kind gift from Dr. Thomas Pucadyil, IISER Pune) using gibson assembly. PCR product was treated with DpnI and transformed into E. coli DH5α bacterial strain for *in-vivo* ligation. Resulting clones were screened by colony PCR. Recombinant protein was expressed in autoinduction medium (Formedium, UK) for 36 hrs at 18°C with constant shaking at 200 RPM.

To obtain human SATB2 expression construct. Coding sequence for human SATB2 was amplified from the IMAGE cDNA clone MGC:119475 IMAGE:40007830 using gene specific primers and were cloned into EcoRI and XbaI digested 3xFLAG-CMV9 vector (Sigma Aldrich, USA). Sequences of all the expression constructs were confirmed using Sanger sequencing and validated for expression using immunoblotting experiments. For each experiment an empty vector was used as control as described in figure legends. Synthetic oligonucleotides used for clonings are obtained from Sigma or Eurofins and are listed below.

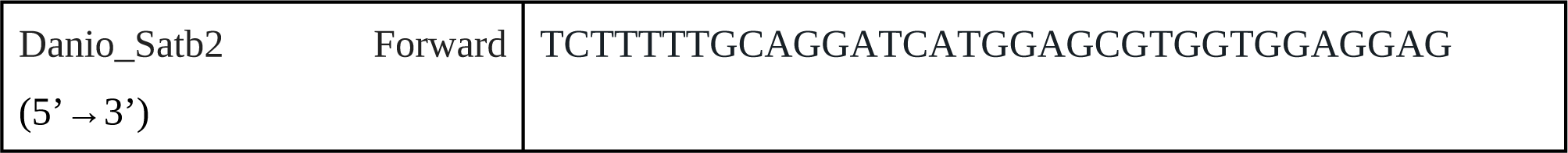
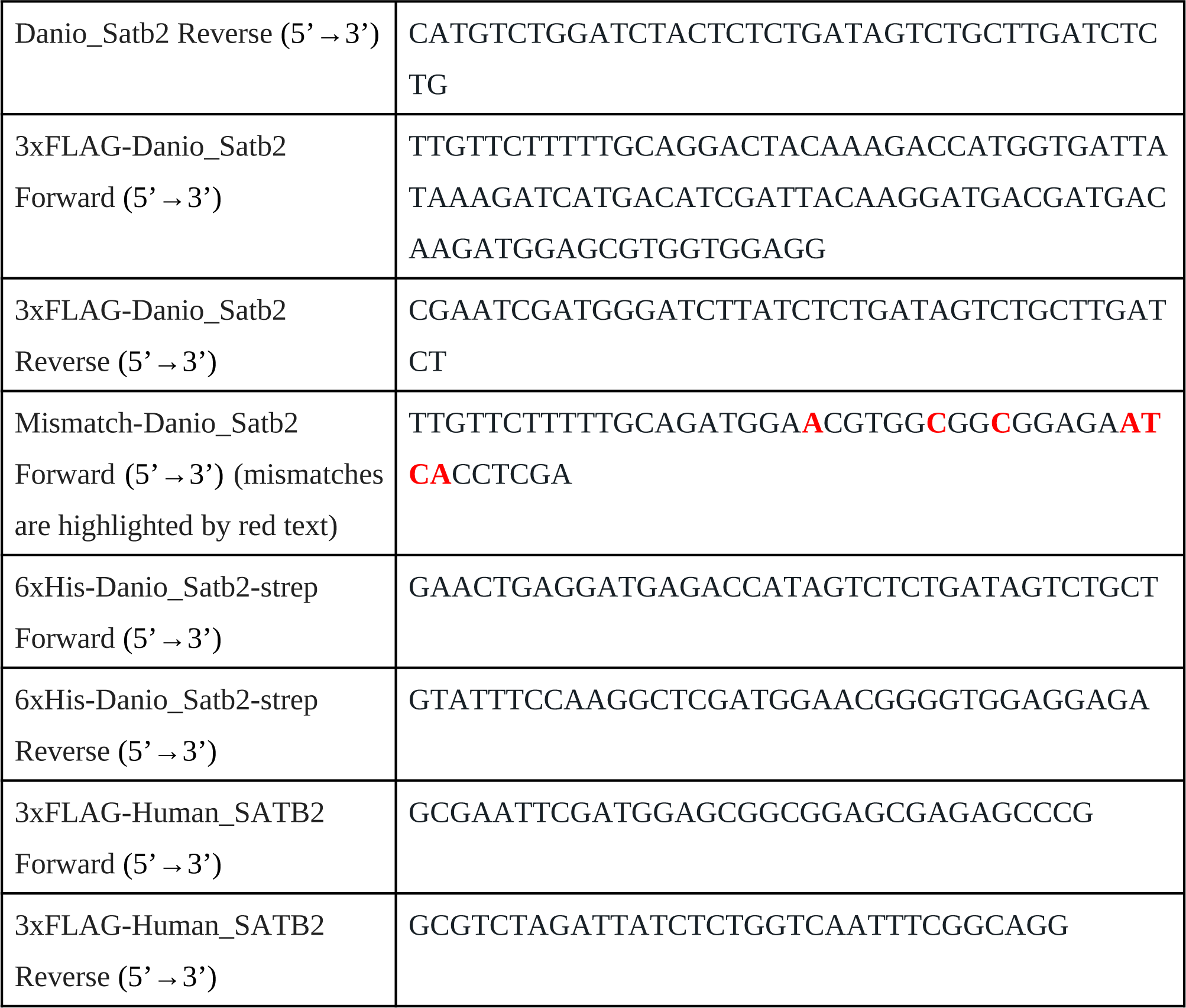

### mRNA and morpholino injections

*In-vitro* mRNA transcription was performed using SP6 mMessage mMachine Kit as per the instructions from the manufacturer (Ambion). Synthesized mRNA was treated with Turbo-DNAse and precipitated using LiCl overnight at −20°C. Synthesized mRNA was checked for quality and quantity by agarose gel electrophoresis and Nanodrop2000 respectively. Small single use aliquots of mRNAs were kept frozen at −20°C till further use. Glass capillaries (WPI) were pulled using a needle puller (P-97, Sutter Instruments) and mounted on a microinjection system (PV820, World Precision Instruments).

For ubiquitous overexpression and morpholino mediated knockdown studies, embryos were arranged in agarose moulds and injections were performed at 1 cell stage as described previously^66^. For overexpression studies, 200pg of either pCS2-Satb2 or pCS2-3XFLAG-Satb2 were injected at 1 cell stage. For single cell gene expression analysis, injections were performed with 200 pg of 3xFLAG-Satb2 at 16 cell stages targeting one cell per embryo to generate mosaicism. For rescue experiments 5pg of morpholino resistant Satb2 version was injected at 1 cell stage followed by injections of 4 ng morpholinos.

The following morpholino sequences targeting translation initiation (MO1) and targeting exon splicing (MO2) were synthesized from Genetools and 1:1 mixture was used for all knockdown experiments. Five base mismatch morpholino was used as a control for all the experiments. Phenotypic changes were captured at indicated time points using Olympus stereo microscope.

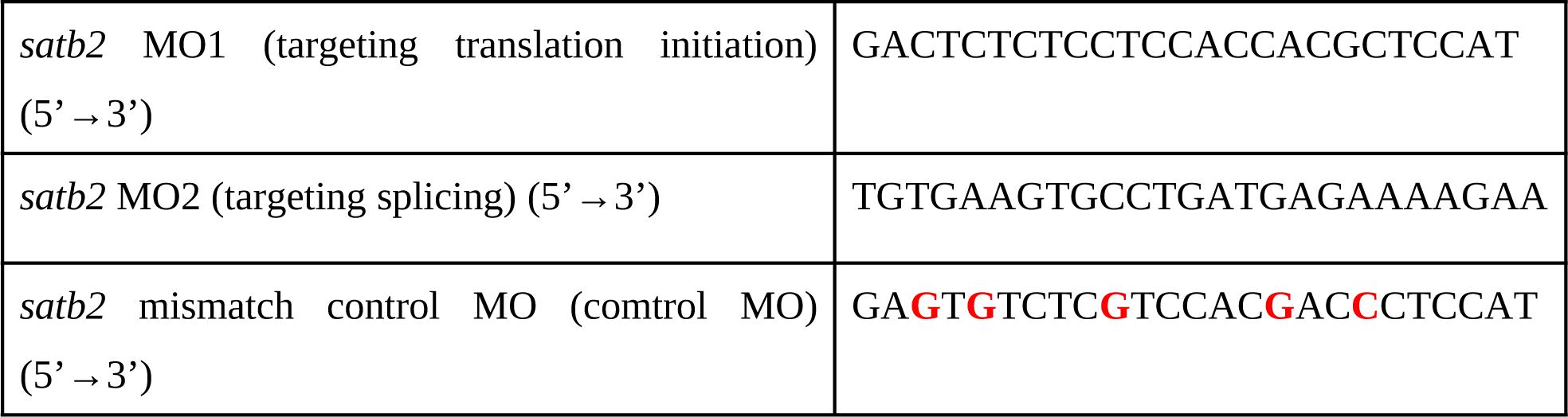

### Genome-wide occupancy analysis for zebrafish Satb2

#### ChIP setup

To map the genome wide occupancy of zebrafish Satb2, we performed ChIP sequencing at desired stages of embryogenesis as mentioned in the results section and figure legends. To validate the use of in-house raised antibody for successful ChIP-seq experiment, we injected embryos with 25pg of 3xFLAG-Satb2 at 1 cell stage and harvested embryos at 4.5 hpf for further processing. ChIP was performed using anti-FLAG antibody (Sigma) and anti-zebrafish Satb2 antibody and subjected to high throughput sequencing as described below. Correlation matrix was generated to confirm the reproducibility of two methods. For all stage specific ChIP experiments, anti-zebrafish Satb2 was used to capture endogenous genomic binding regions.

Briefly, ChIP was performed using modified protocol^5^. Approximately, 2000 embryos per stage were harvested in 1x E3 medium and homogenized using glass homogenizer loose piston in the presence of 1mM PMSF and immediately fixed with 1% methanol free formaldehyde (Thermo) at room temperature for 12 minutes with constant shaking. Fixation was stopped by addition of glycine to a final concentration of 0.125M and incubation at room temperature for additional 5 minutes. Fixed cells were centrifuged at 500g for 5 mins and washed thrice with 1x ice chilled PBS by centrifugation at 4 degrees for 5 minutes each. Cells were further lysed in a 6 times bed volume cell lysis buffer (10mM Tris-HCl pH7.5, 10mM NaCl, 0.5% (v/v) NP-40) for 10 minutes on ice and subjected to homogenization in Dounce homogenizer for 10 times with loose piston followed by 3 times with tight piston. Nuclei were collected by centrifugation at 2500g, washed with ice cold PBS and resuspended in 8 times the pellet volume in the nuclei lysis buffer (50mM Tris-HCl pH7.5, 10mM EDTA, 1% (w/v) SDS, protease inhibitor cocktail (Roche,11873580001)) and incubated further for 30 minutes on ice. The samples were sonicated using the following Covaris S2 sonication conditions: 20% duty cycle, intensity= 5, cycle per burst= 200, Time= 40 cycles of 30 seconds ON 30 seconds OFF to obtain an average size of 200-300 base pairs. The samples were centrifuged at 12000rpm for 10 minutes and the chromatin containing supernatant was stored at −80C till the further use. Prior to ChIP setup, supernatant was precleared using a mixture of 1:1 Dynabeads protein A:G beads for 2 hrs at 4°C. For each ChIP replicate and Mock reaction, 100μg of chromatin was diluted 1:10 with ChIP dilution buffer (16.7mM Tris-HCl pH7.5, 167mM NaCl, 1.2mm EDTA, 0.01% (w/v) SDS, 1xPIC) and incubated with either 5μg of anti-FLAG antibody or 20μg of anti-Satb2 antibody was used. For Mock ChIP reactions, an equal amount of Rabbit IgG (31235 Invitrogen) was used. Chromatin-antibody complex was incubated overnight at 4°C with constant rotating. Following day, Chromatin-Antibody-complex was captured using 100μl pre-blocked (with IgG-free BSA and t-RNA) Dynabeads® Protein A:G mix for 4 hrs at 4°C. Beads were washed with ice-cold buffers in the following order: 7 minutes 4 times with low salt buffer (20mM Tris HCl pH 8.0, 150mM NaCl, 2mM EDTA, 0.1% SDS, 1% Triton X-100), 10 minutes twice with high salt buffer (20mM Tris HCl pH 8.0, 200mM NaCl, 2mM EDTA, 0.1% SDS, 1% Triton X-100), 10 minutes once with LiCl buffer (0.25M LiCl, 1mM EDTA, 10mM Tris HCl pH 8.0, 1% NP40, 1% Sodium deoxycholate) and 10 minutes twice with TE buffer (10mM Tris HCl pH 8.0, 1mM EDTA). The TE buffer was removed completely and 150μL of the elution buffer (0.1M NaHCO3, 1% SDS) was added and vortexed gently to solubilise the beads, followed by incubation at 65C for 30 minutes at 1000rpm. The eluate was collected in a fresh tube and the process was repeated with addition of 150μL of the elution buffer. To 300μL of the eluates and 10% input, 20uL of 5M NaCl and 2μL of RNAseA(10mg/ml) was added and the samples were incubated at 65C overnight with constant shaking at 700-800rpm. Next, 20μL of 1M Tris pH 8.0, 10μL of 0.5M EDTA and 2μL of Proteinase K (20mg/ml) was added and the samples were further incubated at 42°C for 1 hour at 700-800rpm. Samples were purified by standard phenol:chloroform extraction method and DNA was precipitated overnight with equal volume of 100% isopropanol in the presence of Glycoblue (Ambion) at −20°C. DNA pellets were eluted in nuclease free water and quantified using Qubit fluorometer (Thermo) before proceeding with further analysis.

#### Library preparation, sequencing and data analysis

Equal amount of DNA (∼5 ng) was used as an input for library preparation and libraries were prepared using Nugen ultra2 ovation kit (Nugen). Number of cycles for amplification of adapter ligated libraries were estimated by the qPCR method as described in the datasheet provided by the manufacturer. Final libraries were purified using HiPrep PCR clean up system (Magbio). Library concentration was determined using Qubit and average fragment size was estimated using DNA HS assay on bioanalyzer 2100 (Agilent) before pooling libraries at equimolar ratio. Sequencing reads (100bp PE) were obtained on the HiseqX platform at Macrogen Inc, Korea.

Sequencing reads were trimmed using TrimmomaticPE for Truseq2:PE adapters and reads with quality greater than phred 33 were retained^76^. Quality of sequencing reads were determined using fastQC^77^. High quality sequencing reads were aligned to zebrafish danRer10 genome version using default parameters of BWA^78^. Aligned reads were subsampled to 40 million reads in each sample using Bbmap^79^. Correlation between each replicate was estimated using multiBamSummary and replicates showing very high Pearson correlation (> 0.7) were used for further analysis. Peak calling was performed using macs2 with default parameters and q value 0.05. Consensus set of peaks from biological replicates were extracted using a custom R script from Roman Cheplyaka (https://ro-che.info/articles/2018-07-11-chip-seq-consensus). Bigwig files were generated first using bamCoverage normalizing to RPKM and then subtracting Input signals using bamCompare utilities from deepTools 3.3.2^80^ and used for visualization with Integrative Genomics Viewer (IGV). Satb2 peaks in regulatory genes were extracted by intersecting with Ensembl of ATAC seq peaks (GSE106428, GSE130944, GSE101779). Further, Satb2 peaks were classified as putative enhancer bound peaks by intersecting Satb2 peaks with a consolidated dataset of H3K4me1 (GSE32483, GSE74231). Promoter bound peaks were assigned as +/− 2 kb from the TSS of the genes. Peak annotation to the nearest gene was performed using annotate.pl utility from Homer. Motif discovery for a given set of peaks were performed using findMotifsGenome.pl script from Homer and gene ontology was performed using webGestalt utilizing KEGG and Panther databases.

### ChIP qPCR

ChIP experiment was performed as described above at desired stages in replicates as indicated in figure legends. Equal amount of IPed DNA was used as an input for quantitative real time PCR analysis. qPCR was performed using primers listed below with KAPA Sybr green master mix (Kapa biosystems) on ViiA7 Real-time pcr system (ABI). Relative enrichment was calculated using percent input method using formula 100 × 2~(Adjusted input - Ct (IP).

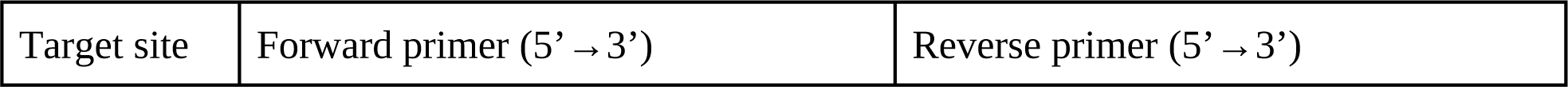
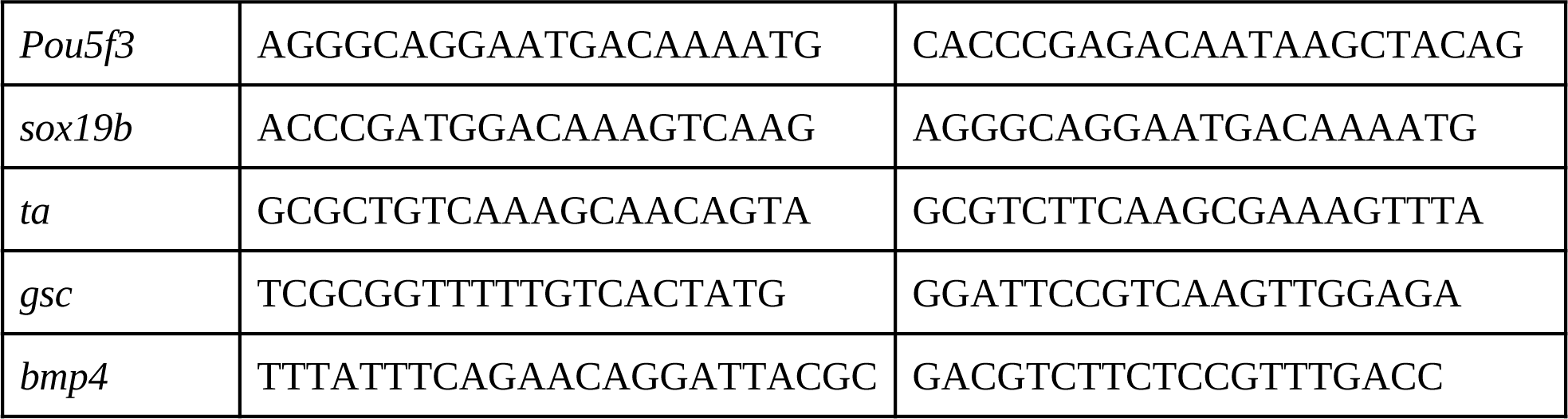

### Genome-wide occupancy analysis for mouse SATB2

#### ChIP setup

To obtain genome wide binding sites for SATB2 during early neurogenesis, SWR/J mice were incrossed and embryos were isolated at desired stage of development. For experiments with E9.5 head tissue and trunk (visceral organs were discarded) were dissected under stereo microscopes with fine forceps and collected in ice chilled 1xPBS containing 0.5% glucose. Tissues from 10-12 embryos were pooled together for each replicate and crosslinked using 1% methanol free formaldehyde for 8 minutes followed by quenching with 125 mM glycine for additional 5 minutes at room temperature. For experiments at E13.5, dorsal telencephalon was isolated and processed as described above. Chromatin isolation and ChIP experiment was performed as described in experiments for zebrafish Satb2 with modifications as follows. 100 µg of precleared chromatin was used for each IP reaction and 10 µg of anti-Human SATB2 antibody was used for pulldown. Equal amount of anti-Rabbit IgG (Invitrogen) was used as a control for each ChIP experiment. Antibody-protein complex was captured using 100 µl of preblocked Dynabeads protein A:G mixture for 4 hours at 4°c. Samples were processed further as described in the earlier section.

#### Library preparation, sequencing and data analysis

Equal amount of DNA (∼5 ng) was used as an input for library preparation and libraries were prepared using NEB ultra II DNA library prep kit (NEB). Sequencing reads (76 bp PE) were obtained on the Nextseq 550 platform at IISER, Pune.

Sequencing reads were trimmed using TrimmomaticPE for Truseq2:PE adapters and were aligned to mouse mm10 genome using default parameters of BWA. Aligned reads were subsampled to 25 million reads for each sample using BBmap. QC, peak calling and bigwig generation was performed as described in the above sections.

### ChIP Western

Efficiency of ChIP for mouse SATB2 and validation of mammalian SATB2 antibody was performed by ChIP western assay. Briefly, ChIP was performed with in house anti-human Satb2 antibodies. After bead washing and removal of excess TE buffer, beads were resuspended in 1x PBS and treated with 2µl of DNAse (10mg/ml) for 30 mins at 37°C. Further, beads were washed with 1x PBS twice and boiled in 2X laemmli buffer (0.25M Tris-Cl pH 6.8, 1% SDS, 1% β-mercaptoethanol, 15% glycerol) at 98°C for 5 minutes and electrophoresed on 7.5% of SDS-PAGE gel. Proteins were transferred to the 0.45µM PVDF membrane (millipore) by wet transfer method at 0.6A for 2.5 hrs at 4°C. Non-specific sites on the membrane were blocked using 3% BSA and further incubated with anti-SATB2 antibody (1:1000 abcam ab34735) in 0.3% BSA overnight at 4°C with constant rocking. Next day, Membrane was washed with 1X TBST (50mM Tris-Cl pH 7.4, 150mM NaCl, 0.1% Tween-20) 4 times for 7 minutes each and incubated with anti-Rabbit HRP conjugated antibody (1:10000 STAR124P Bio Rad) in 1x TBST for 45 minutes at room temperature. After removing excess of secondary antibody by repeated washes with 1x TBST, signal was developed with Clarity western ECL substrate and captured with LAS 4000 system (GE healthcare).

### ChIP sequencing for histone modification marks

#### ChIP setup

To profile the status of histone modifications upon over expression of 3xFLAG tagged Satb2, 1000 embryos were harvested for each ChIP reaction at 4.5 hpf (Dome stage) in batches until sufficient amount of input material is generated. Empty pCS2-GFP was used as a control for injections. Pulldowns were performed with 20 µg chromatin and 5 µg anti-H3K4me3 (ab8580 abcam), 5 µg anti-H3K27Ac (ab4729 abcam), 5 µg anti-H3K27me3 (07-449 Millipore) and 5 µg of normal rabbit IgG overnight at 4°C. Antibody-Protein complex was captured using 50 µl of Preblocked Dynabeads A:G mixture for 3 hrs at 4°C. Samples were processed further as described in the earlier section.

#### Library preparation, sequencing and data analysis

5ng of purified ChIP DNA was used for library preparation using NEB ultra II DNA kit (NEB) as described in the earlier section. Sequencing reads (100bp PE) were obtained on the HighseqX platform at Macrogen Inc, Korea. Sequencing reads were trimmed using TrimmomaticPE for Truseq2:PE adapters and reads with quality greater than phred 33 were retained^76^. Quality of sequencing reads were determined using fastQC^77^. High quality sequencing reads were aligned to zebrafish danRer10 genome version using default parameters of BWA^78^. Aligned reads were subsampled to 40 million reads in each sample using Bbmap^79^. RPKM normalised Input signal subtracted BigWig tracks for visualization were generated using bamCompare tool from deepTools 3.3.2.

### ATAC-seq and data analysis

Single embryos at 14 somite stages from the incross of *satb2*^+/−^ were hand dechorinated and harvested in 50 ul of ice chilled 1x DPBS (Invitrogen). 5 µl of cell suspension was used for identifying genotypes of each embryo. Cell suspension from 3 embryos were pooled together and processed for Omni-ATAC seq as described previously^81^ with modification from Amanda Ackermann lab. Briefly, cells were washed with 1x DPBS and resuspended in cell lysis buffer (10 mM Tris pH 7.5, 10 mM NaCl, 3 mM KCl, 0.1 % NP40, 0.1% Tween20 and 0.01% Digitonin) and incubated on ice for 3 minutes. Further, cells were washed with a wash buffer (10 mM Tris pH 7.5, 10 mM NaCl, 3 mM KCl and 0.1% Tween20) by centrifugation at 500 for 10 mins 4°C. Supernatant was discarded and pellet was resuspended in 25 µl 2x Tagmentation buffer (Illumina, catalog # 15027866), 16.5 µl DPBS, 0.5 µl 10% Tween 20, 0.5 µl 1% Digitonin, 5 µl nuclease and 2.5 µl Tn5 transposase enzyme (TDE1, Illumina, catalog # 15027865) and incubated for 28 minutes at 37°C. After the tagmentation reaction, DNA was isolated using the Qiagen MinElute Reaction Cleanup Kit (Qiagen). Purified DNA was used as an input to generate a library by amplifying with 2x Q5 DNA polymerase mix (NEB) and indexing primers. Optimal cycles were determined using qPCR analysis. Amplified libraries were purified using Agencourt ampure XP beads to remove adapters and larger fragments.

For Satb2 overexpression studies, 200 pg 3xFLAG Satb2 were injected at 1 cell stage. Embryos (a pool of 10 embryos for each replicate) were harvested at 4.5 hpf and processed for ATACseq as described above.

Sequencing reads (41 bp PE) were obtained on NExtseq 550 at IISER Pune and trimmed for Nextera adapters using default parameters of Trimmomatic PE. Trimmed reads were aligned to danRer10 using default parameters of Bowtie2^82^. Briefly, BAM files were subsampled to 55 million reads in each sample using bbmap and sorted by name. paired end bed files were obtained using bedtools bamtobed. Reads were displaced by +4 bp and -5 bp. Peak calling was performed using macs2 callpeak -f BEDPE -q 0.05 -- nomodel --extsize 200 --gsize 1.3e9 --keep-dup 2 parameters. Consensus peaks were obtained using a custom R script used for ChIPseq analysis. BigWig files were generated using bamCoverage (deepTools). Peaks were annotated to the nearest gene using Homer and classified into promoter (+/− 2 kb) and non-promoter regions. K-means clustering was performed around +/− 2kb of Satb2 peak center using deepTools. Clusters were annotated using Homer and gene ontology analysis was performed using webGestalt. Motif analysis was performed using findMotifsGenome.pl from Homer.

### Differential chromatin accessibility analysis (Diffbind)

The differential chromatin accessibility analysis of Satb2 mutant and wild type embryos at 14 somite stages was performed using the DiffBind^83^. Significantly differentially accessible peaks were identified using the Deseq2 package and only sites with FDR < 0.05 and fold change of > Log2 (+/− 1.5) were used for further analysis. Differential accessible sites were annotated to the nearest gene using Homer. Core promoter was defined as +/− 2 kb from the TSS.

### Nucleosome occupancy analysis

Nucleosome occupancy analysis was carried out using the nucleoATAC suite with default parameters for Satb2 mutant and wild type ATACseq datasets. Nucleosome fuzziness scores were obtained and used for calculating the difference in nucleosome phasing upon loss of function of Satb2.

### Single cell RNAseq analysis

To generate single-cell gene expression datasets, a mosaic over expression system was generated by injecting one cell of 16 cell stage embryos with 200 pg of 3xFLAG-Satb2. Only one cell per embryo was injected randomly irrespective of its spatial arrangement. Embryos were raised at 28.5°C, hand dechorinated and dissociated further in 1x DMEM-F12 + 125 mM EGTA by hand tapping. Cells were collected by centrifugation at 300 g for 2 minutes and resuspended in 1X DPBS + 0.1% BSA. cell suspension was passed through a 70-μm Flowmi cell strainers (Sigma). Cell viability was estimated using CountessII automated cell counters and only samples with more than 95% viability were processed further with 10x Genomics Chromium. Approximately 8000 cells were captured in GEMs. scRNA-seq libraries were prepared as described in 10x Genomics manuals (Single Cell 3′ Reagent Kits V3, User Guide PN-1000092). Immediately following GEM generation, reverse transcription reaction was carried out using eppendorf mastercycler pro by incubating at 53°C for 45 minutes followed by denaturation at 85°C for 5 minutes. Single-stranded cDNA was purified using DynaBeads MyOne Silane Beads (Thermo). cDNA amplification was performed for 11 cycles with initial denaturation at 98oC 3 minutes followed by repeated cycles of 98oC for 15 seconds, 63oC for 20 seconds and 72oC for 1 minute. Final extension was performed at 72oC for 1 min. cDNA quality was determined on a Bioanalyzer 2100 high sensitivity assay. cDNA was further fragmented, end repaired and A-tailed according to manufacturer’s instructions. Before proceeding for adapter ligation, samples were purified using double-side clean up protocol with SPRI bead (Beckman Coulter). Adapter ligated sample was subjected to amplification for 11 cycles using indexing primer from Chromium i7 Multiplex Kit (PN-120262). Amplified libraries were again subjected to double-side purification using SPRI and quantified using Qubit fluorometer and library size was estimated using Bioanalyzer 2100.

1.5 pM of the denatured library was used as an input to obtain sequencing reads using 28 cycles for read1, 8 cycles for indexes and 101 cycles for read2 on Nextseq 550 at IISER Pune.

Sequencing data was further processed with default parameters using cell Ranger 3.0.2 (10X Genomics). Sequencing reads were aligned to danRer10 genome using STAR aligner. Feature matrices generated using cell Ranger were utilized for further analysis using Seurat 3.0.

### TF binding promoter scan

Consensus binding sites for respective TFs were extracted from JASPAR database and MAST (MEME Suite) was used for scanning genomic regions extracted from UCSC server using version danRer10.

### NT2/D1 cell culture

Human embryonic carcinoma cell line NT2/D1 (NTERA2-clone D1) were a kind gift from Dr. Peter Andrews, University of Sheffield, UK. They were cultured in Dulbecco’s Modified Eagle’s Medium with sodium pyruvate, high glucose (DMEM, Sigma-Aldrich, St. Louis, Missouri, USA) supplemented with 10% foetal bovine serum (Invitrogen, Carlsbad, California, USA), 2 mM L-glutamine (Invitrogen, Carlsbad, California, USA) and penicillin-streptomycin (Invitrogen, Carlsbad, California, USA) and maintained at 37°C under 5% CO2 atmosphere. NT2/D1 cells were subcultured upon reaching 70% confluency by gentle scraping.

### Transfection of overexpression constructs in NT2/D1

For overexpression of SATB2 in NT2/D1 cells, 0.8 × 10^6^ cells were seeded in 60mm cell culture grade plates. 16 to 20 hours post-seeding, the cells were transfected with either empty FLAG vector or FLAG-SATB2 constructs for overexpression using Lipofectamine 2000 (Invitrogen, Carlsbad, California, USA) as per manufacturer’s guidelines.

### All-trans retinoic acid mediated differentiation of NT2/D1

All-trans-retinoic acid (RA) (Sigma-Aldrich, St. Louis, Missouri, USA) was reconstituted at a concentration of 5 mg/ml in DMSO (Sigma-Aldrich, St. Louis, Missouri, USA) and stored at −80°C. For differentiation, NT2/D1 cells were harvested using 0.05% trypsin resuspended in fresh medium and seeded at a density 1 × 10^6^ cells in 100 mm tissue culture dish (Corning, New York, USA). Cells were allowed to grow for 24 hours following which 1 × 10^7^ cells were harvested for RNA and protein extractions as 0 day control. RA was added to the remaining plates at a concentration of 13.7 μM for the rest of 5 days. Each day cells were either replenished with fresh medium and RA or harvested for RNA.

### qPCR analysis

Embryos were collected at the desired stage and were lysed in RNA iso plus (DSS-Takara, India) to extract total RNA. Each sample consisted of at least 10 embryos per experiment, and all experiments were repeated at least three times independently. cDNA was synthesized using High capacity cDNA synthesis kit (Applied Biosystems), following manufacturer’s instruction.

Quantitative real time PCR was performed using SYBR green chemistry (KAPA biosystems) using gene specific primers as listed below on ViiA7 thermal cycler (Applied Biosystems). Changes in threshold cycles were calculated by subtracting the Ct values of the gene of interest from that of housekeeping control (for qRT-PCR) [Ct(target genes) – Ct(*ef1a* or GAPDH)]. ΔCt values of specific target genes from the experimental samples were then subtracted from their respective control samples to generate ΔΔCt values. The fold changes were calculated using the formula : 2~(−(ΔΔCt value).

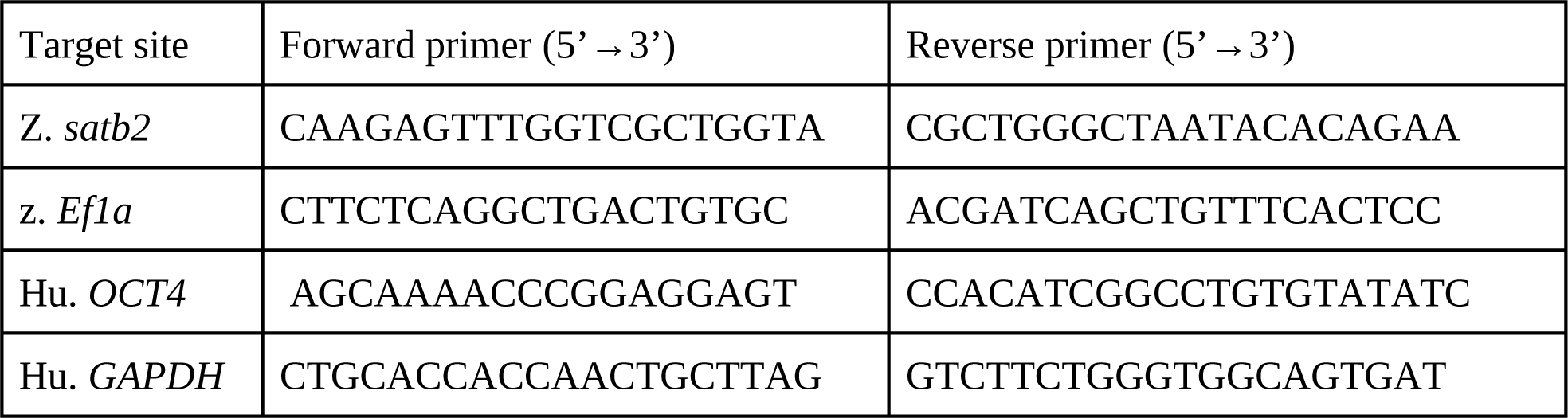

### Protein extraction and immunoblotting

To confirm the absence of Satb2 in zebrafish larvae. Larvae were manually dechorionated, deyolked and harvested at 48 hpf by boiling in 2X laemmli buffer (0.25M Tris-Cl pH 6.8, 1% SDS, 1% β-mercaptoethanol, 15% glycerol) at 98°C for 5 minutes and electrophoresed on 7.5% of SDS-PAGE gel. Proteins were transferred to the 0.45µM PVDF membrane (millipore) by wet transfer method at 0.6A for 2.5 hrs at 4°C. Non-specific sites on the membrane were blocked using 3% BSA and further incubated with inhouse anti-SATB2 antibody (1:500 This study) in 0.3% BSA overnight at 4°C with constant rocking. Next day, Membrane was washed with 1X TBST (50mM Tris-Cl pH 7.4, 150mM NaCl, 0.1% Tween-20) 4 times for 7 minutes each and incubated with anti-Rabbit HRP conjugated antibody (1:10000 STAR124P Bio Rad) in 1x TBST for 45 minutes at room temperature. After removing excess of secondary antibody by repeated washes with 1x TBST, signal was developed with Clarity western ECL substrate and captured with LAS 4000 system (GE healthcare).

For experiments with NT2D1 Cells pellets were resuspended in RIPA buffer (10 mM Tris (pH 8.0), 1 mM EDTA (pH 8.0), 0.5 mM EGTA, 1% Triton X-100, 0.1% sodium deoxycholate, 0.1% SDS, 140 mM NaCl) containing 1X protease inhibitors (procured from Roche, Basel, Switzerland) and lysed by repeated freeze-thaw cycles. The lysates were centrifuged at 14000 rpm, 4°C, 30 minutes to eliminate the cellular debris. The supernatant was collected in the fresh microfuge tube. The concentrations of protein were determined by performing BCA assay (purchased from ThermoFisher Scientific, Waltham, MA, USA). Equal amounts of protein lysates were boiled in 1X Laemmli buffer (0.5 M Tris-HCl pH 6.8, 28% glycerol, 9% SDS, 5% 2-mercaptoethanol, 0.01% bromophenol blue) for 10-15 minutes and subjected to electrophoresis on a polyacrylamide gel. The separated proteins were transferred onto PVDF membrane (Millipore, Billerica, Massachusetts, USA) using phosphate-based transfer buffer (10 mM sodium phosphate monobasic, 10 mM sodium phosphate dibasic) at 4°C, 600 mA, 2 hours. After the completion of transfer, membranes were blocked in 5% skimmed milk, incubated overnight at 4°C with the anti-FLAG antibody prepared in 5% BSA. The membranes were washed thrice with the buffer containing 20 mM Tris buffer pH 7.4, 500 mM NaCl and 0.1% tween 20 (TST) the next day and incubated with the appropriate secondary antibodies conjugated with horseradish peroxidase for an hour at room temperature. Following this, the membranes were again washed thrice with TST buffer. The blots were developed using Immobilon Western Chemiluminescent HRP Substrate (Millipore, Billerica, MA, USA) and detected using ImageQuant LAS 4000 (GE Healthcare, Piscataway, NJ, USA) according to the manufacturer’s instructions.

### Statistical analysis

GraphPad Prism (GraphPad software, USA) was used for statistical analysis. The type of analysis used in each experiment is mentioned in the respective figure legends.

## Supplemental Figures

**Supplementary Fig. 1.**
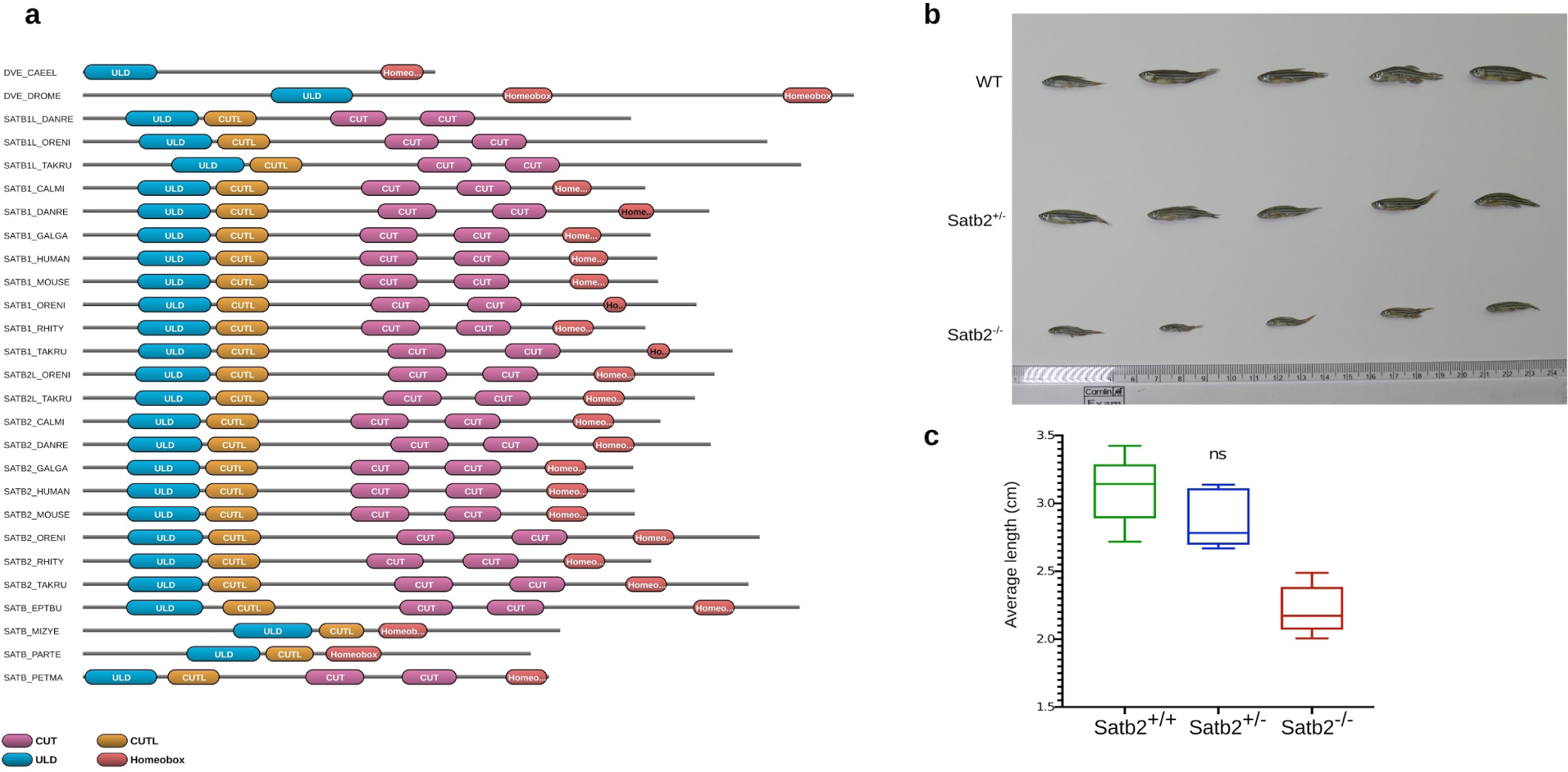
Domain architecture analysis of SATB proteins and characterization of Satb2 mutants. **a**, Comparative analysis of domain architecture of SATB1 and SATB2 for various species across evolution. ULD domain is represented in blue, CUT like domain in yellow, CUT domains in purple and Homeobox domain in red. Homeobox domain with black text indicates putative or predictive Homeobox. **b**, Digital snapshots of 4 months old zebrafish Satb2 mutants along with wild-type and heterozygous siblings highlighting defects in body growth. **c**, Box plot indicating the difference in body length between Satb2 mutants and siblings (n=5, ** p-value <0.05).

**Supplementary Fig. 2.**
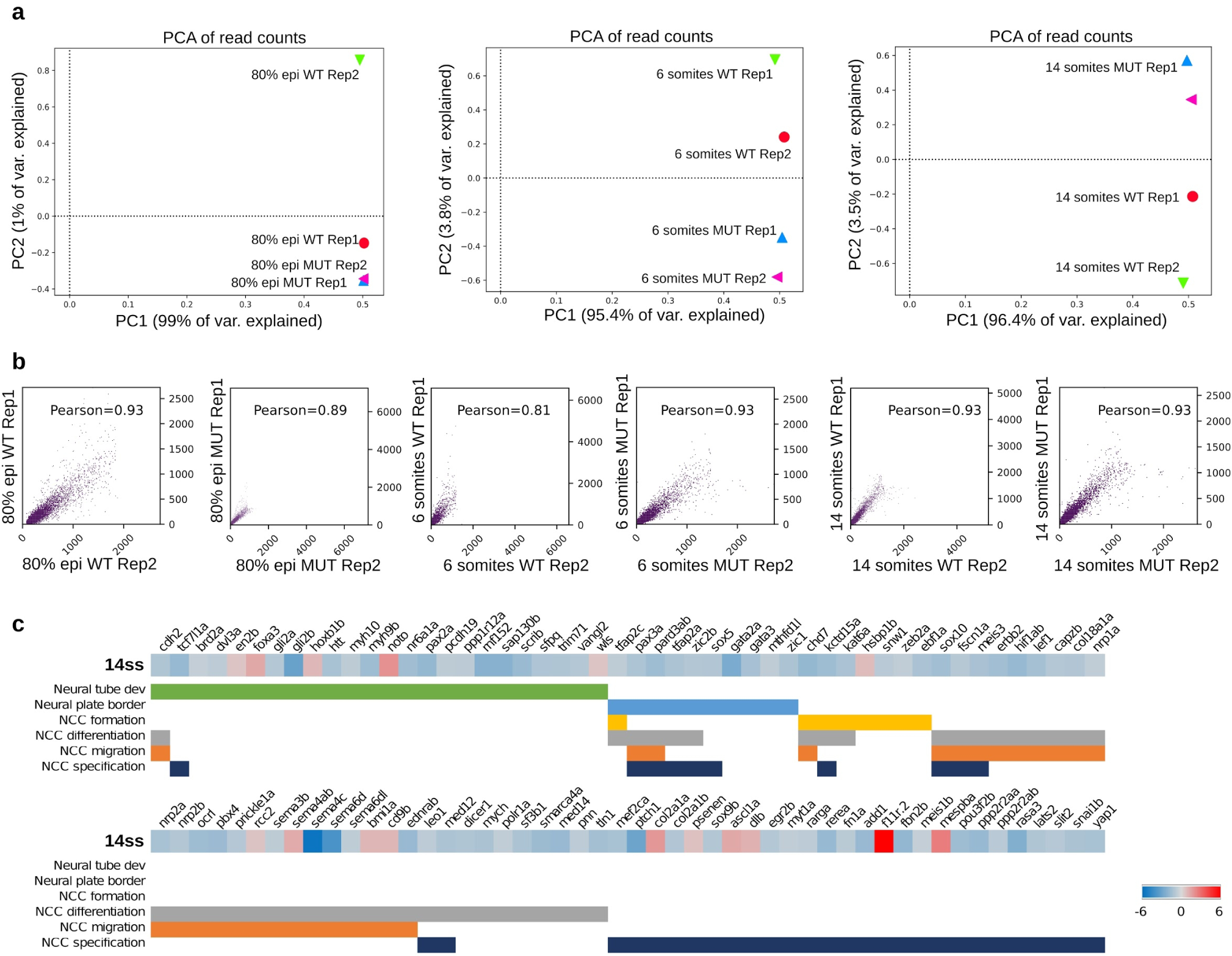
Correlation analysis between replicates of 3’ mRNA seq experiments confirming reproducibility and classification of DE genes at 14 somites. **a**, PCA plots showing highlighting reproducibility between the replicates of 3’ mRNA seq at 80% epiboly, 6 ss and 14 ss respectively. **b**, Pearson correlation analysis between replicates of transcriptome samples at the corresponding stages. **c**, Heatmap of DE genes in 14 ss mutants classified according to their known function during neurogenesis. Genes involved in neural tube development are marked using a green bar, genes involved in neural plate border marked in blue, in NCC formation by yellow, in NCC differentiation by grey, in NCC migration by orange and in NCC specification by dark blue. Colour bar indicates the scale of log2 fold change expression values compared to wild-type.

**Supplementary Fig. 3.**
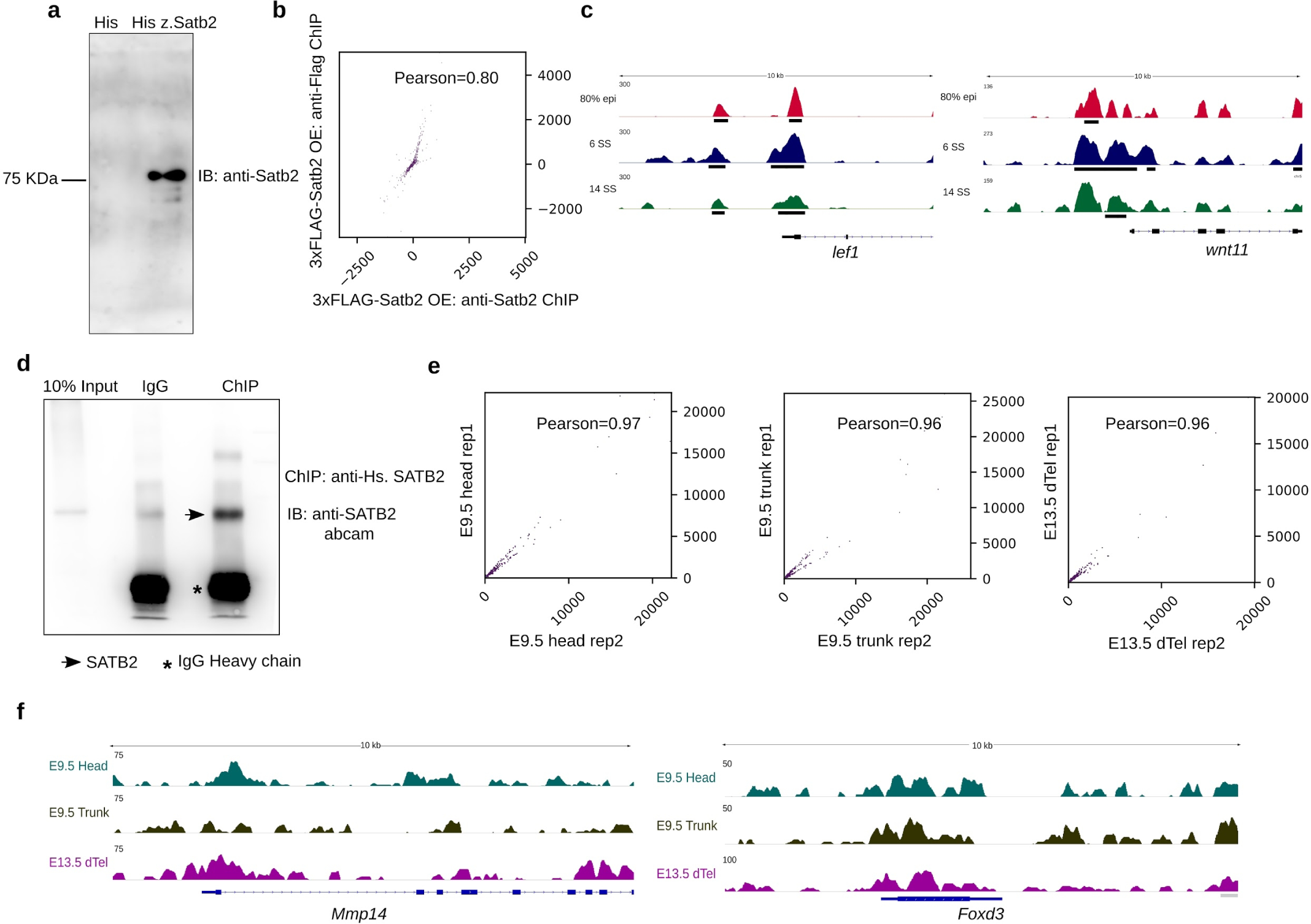
Characterization of anti-SATB2 antibodies and Correlation analysis of zebrafish Satb2 and mouse *SATB2* ChIP seq experiments. **a**, Immunoblot analysis of recombinant control His tag and His-tagged zebrafish Satb2 protein using lab generated zebrafish specific anti-Satb2 antibody. **b**, Pearson correlation analysis of ChIP-sequencing datasets obtained using anti-FLAG antibody and anti-SATB2 antibody showing a high level of correlation. **c**, Integrative genome viewer snapshots of Satb2 occupancy on genomic loci of zebrafish *lef1* and *wnt11* genes. **d**, ChIP western analysis to validate the efficiency of mammalian anti-SATB2 specific antibody. **e**, Pearson correlation plots highlighting reproducibility between replicates of mousse ChIP-sequencing samples at corresponding stages and tissues. **f**, Integrative genome viewer snapshots of SATB2 occupancy on genomic loci of mouse *Mmp14* and *Foxd3* genes.

**Supplementary Fig. 4.**
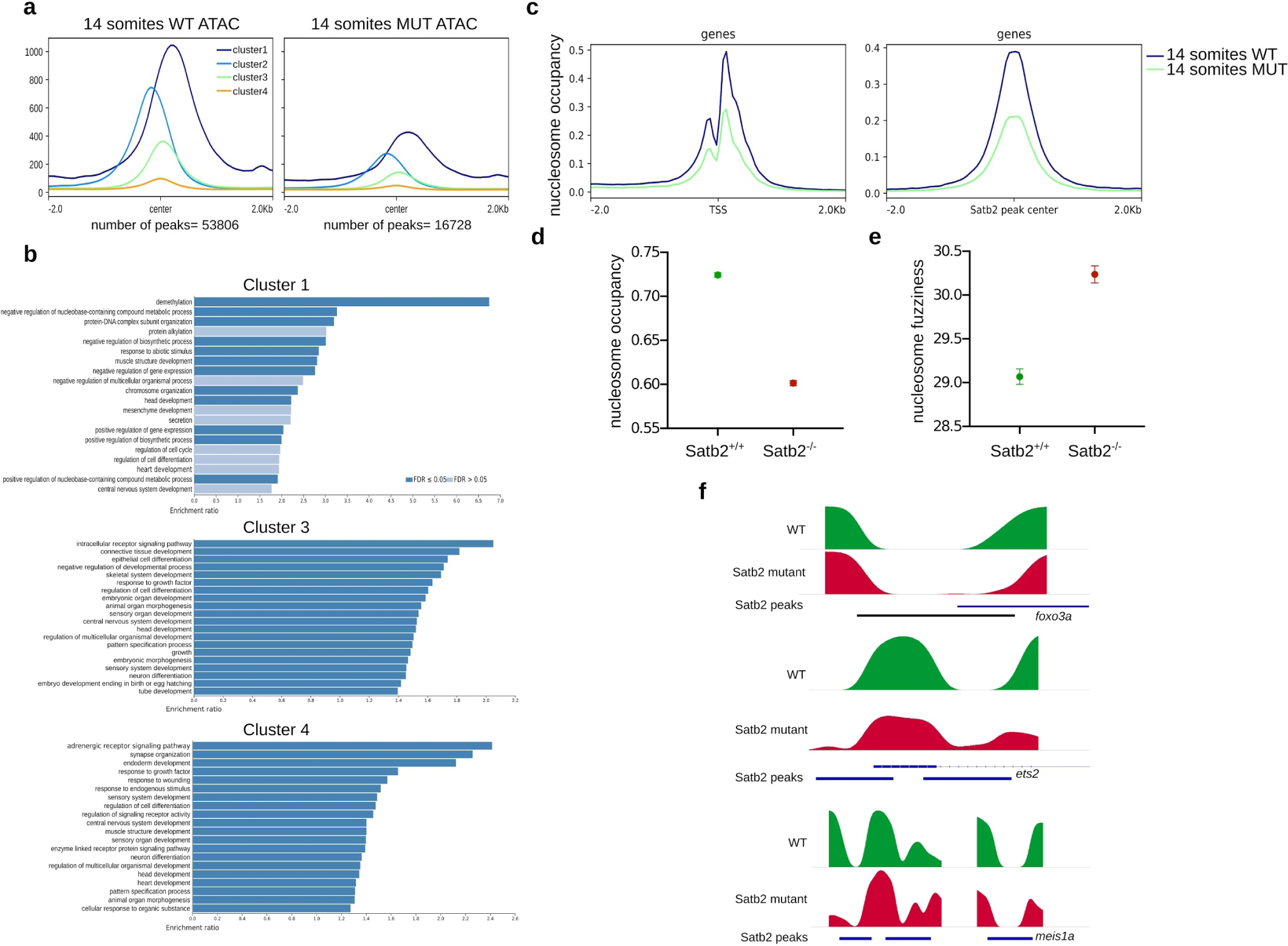
Characterization of chromatin accessibility in Satb2 mutants compared to wild-type siblings at 14 somites. **a**, K-means clustering of average profiles of chromatin accessibility for wild type and satb2^-/-^ at 14 ss. **b**, Gene ontology analysis for cluster1, cluster3 and cluster4 for K-means clustering of chromatin accessibility. **c**, Mean density maps for average nucleosome occupancy profiles in wild type and Satb2 mutants centred around TSS and Satb2 ChIP peak centre respectively. **d**, Dot plot depicting a decrease in average nucleosome occupancy in Satb2 mutants as compared to wild type (**** p-value < 0.0001). Error bar indicates +/− SEM. **e**, Dot plot depicting an increase in average nucleosome fuzziness in Satb2 mutants as compared to wild type (**** p-value < 0.0001). Error bar indicates +/− SEM. **f**, Integrative genome viewer snapshots of nucleosome occupancy tracks on the genomic loci of upregulated genes *foxo3a*, *ets2* and *meis1a* upon loss of Satb2.

**Supplementary Fig. 5.**
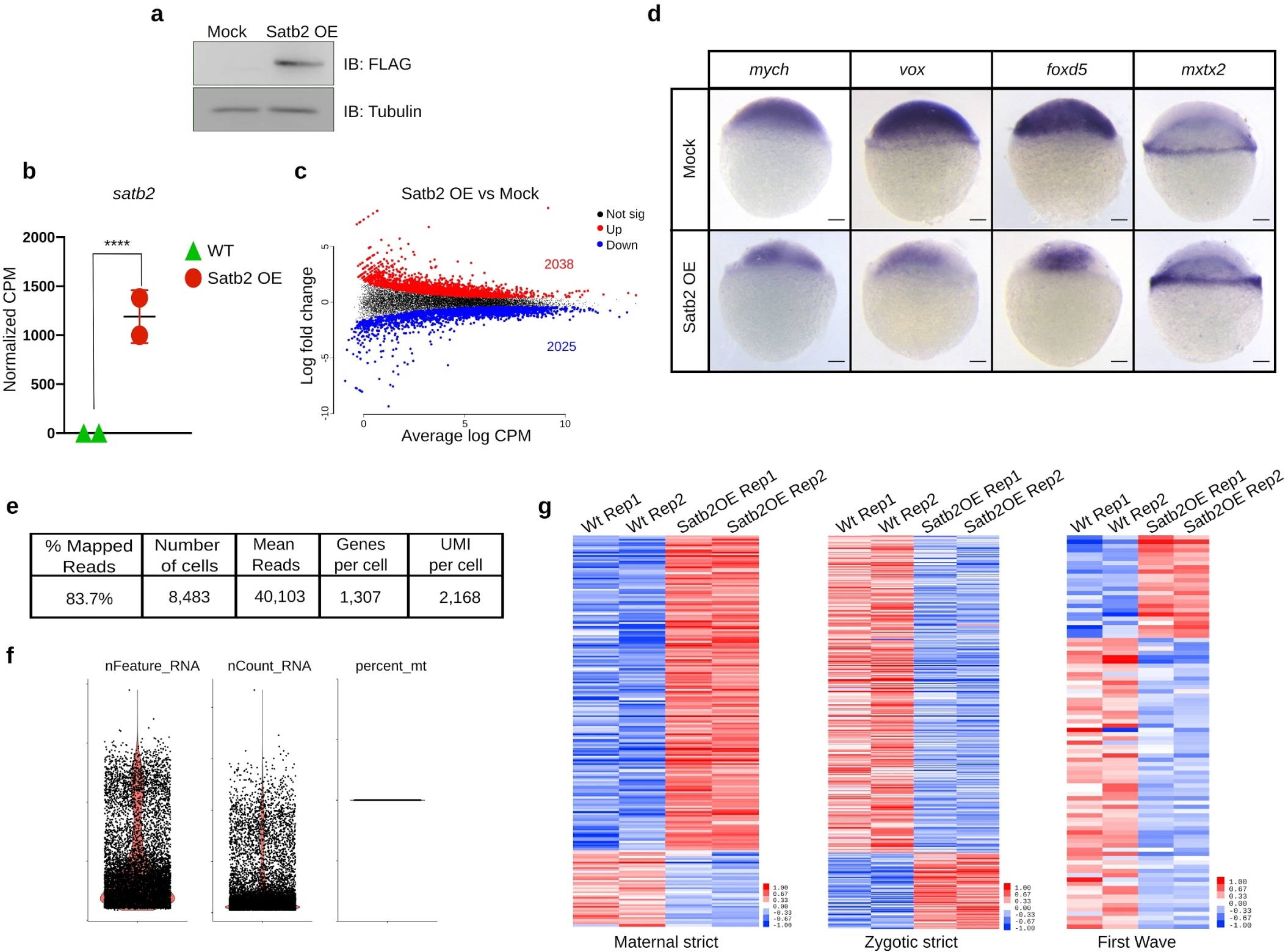
Characterization of the gain of function of maternal Satb2. **a**, Immunoblot analysis for validating overexpression of 3x-FLAG Satb2 at dome stage using anti-FLAG antibody. gamma-Tubulin was used as a loading control. **b**, Dot plot representing and confirming overexpression of Satb2 from RNAseq studies performed at dome stage (4.5 hpf). Normalized CPM counts were used for the analysis (**** p-value 0.0001). **c**, Smear plot analysis for bulk mRNA seq analysis in Mock and 3xFLAG-Satb2 injected embryos at 4.5 hpf. Values in red and blue represent the number of genes differentially upregulated and downregulated respectively. **d**, Whole-mount in situ mRNA analysis showing the effect of Satb2 overexpression on expression of *mych*, *vox, foxd5* (downregulated genes) *mxtx2* (upregulated upon overexpression). **e**, Table representing statistics associated with single-cell RNA seq experiment. **f**, Violin plot for visualization of QC metrics of single-cell transcriptome study. **g**, Heatmaps representing differential expression patterns for strictly maternally expressed genes, strictly zygotically expressed genes and first wave zygotic genes.

**Supplementary Fig. 6.**
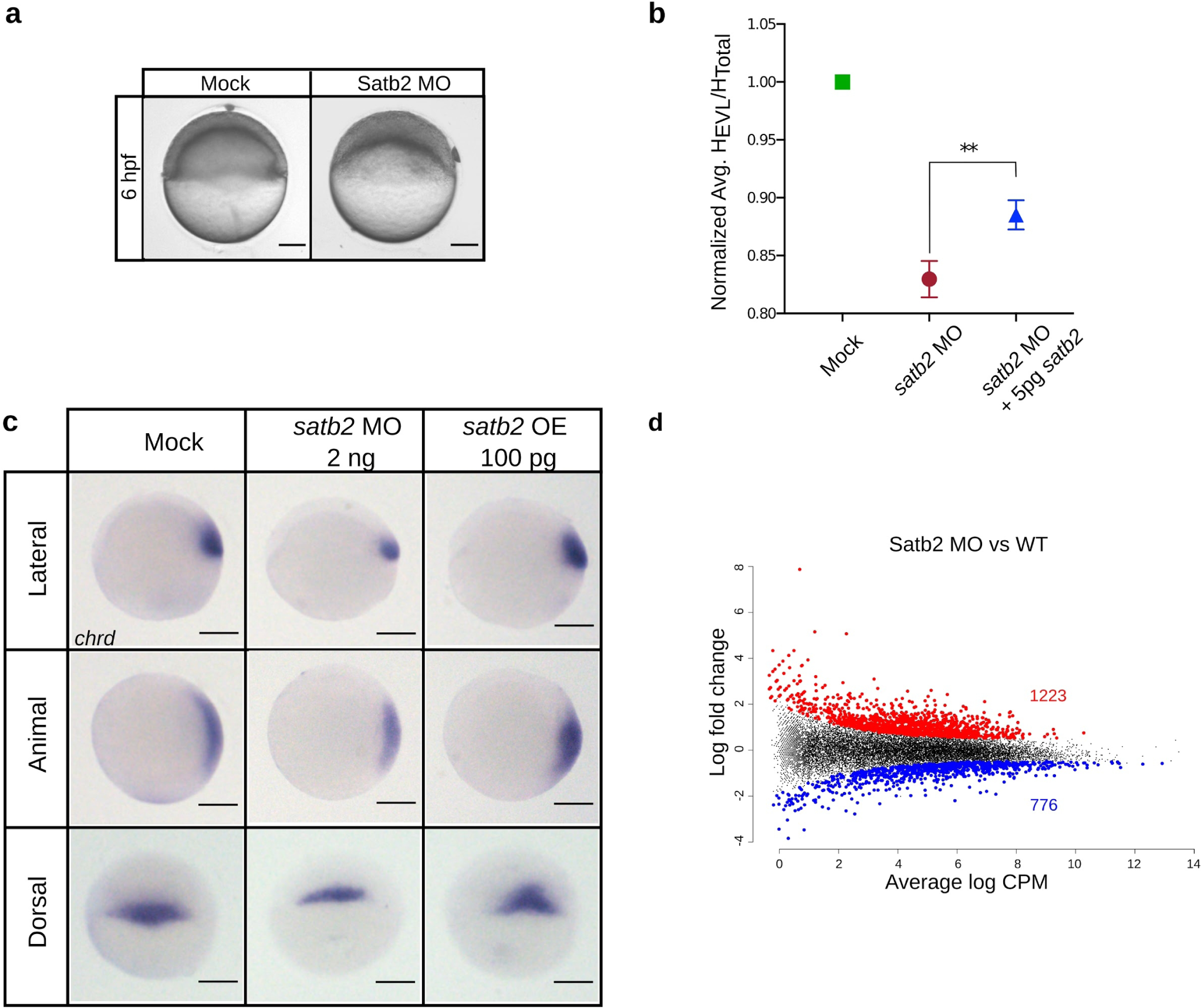
Characterization of the loss of function of maternal Satb2. **a**, Lateral view of zebrafish embryos at 6 hpf injected with 4 ng of control and Satb2 targeting morpholinos. **b**, Dot plot representing ratio of average height of EVL from animal pole to the total height of the embryo normalized to Mock control (n=30). ** signifies P value < 0.005 as determined by student’s t-test. **c**, Whole-mount in situ mRNA analysis for stage matched embryos showing the effect of Satb2 knockdown and over expression on the expression of dorsal marker *chrd*. **d**, Smear plot analysis for bulk mRNA seq analysis in Control and Satb2 morpholino injected embryos at 4.5 hpf. Values in red and blue represent the number of genes differentially upregulated and downregulated respectively.

**Supplementary Fig. 7.**
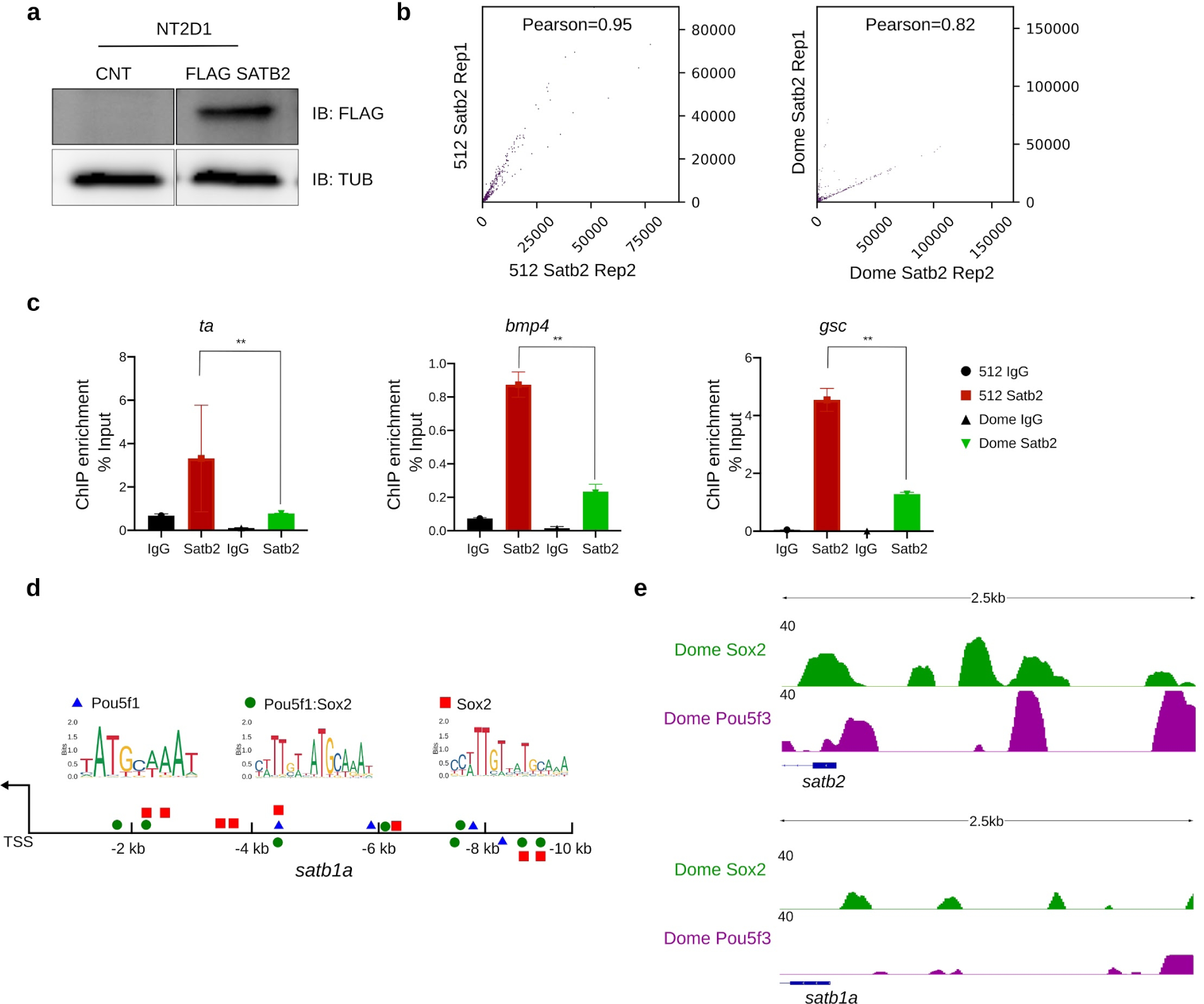
Interplay between Satb2 and pluripotency factors. **a**, Immunoblot analysis for validating overexpression of 3x-FLAG SATB2 in NT2D1 cells using anti-FLAG antibody. gamma-Tubulin was used as a loading control. **b**, Pearson correlation analysis of ChIP-sequencing datasets obtained using anti-Satb2 antibody at 512 cells stage and at the dome stage showing a high level of correlation. **c**, Quantitative relative enrichment (ChIP) represented as percent input using isotype-matched IgG and anti-Satb2 antibody for genomic locus (TSS) of negatively regulated genes *ta*, *bmp4* and *gsc*. ** indicates P-value < 0.001 as calculated by the Student’s two-tailed t-test, N=2. **d**, Schematic for related protein *satb1a* promoter displaying motif sites marked for Pouf53, Pou5f3: Sox2 and Sox2. **e**, Integrative genome viewer snapshots of Pou5f3 occupancy on genomic loci of zebrafish *satb2* and *satb1a* respectively highlighting selective regulation by Pou5f3.

**Supplementary Fig. 8.**
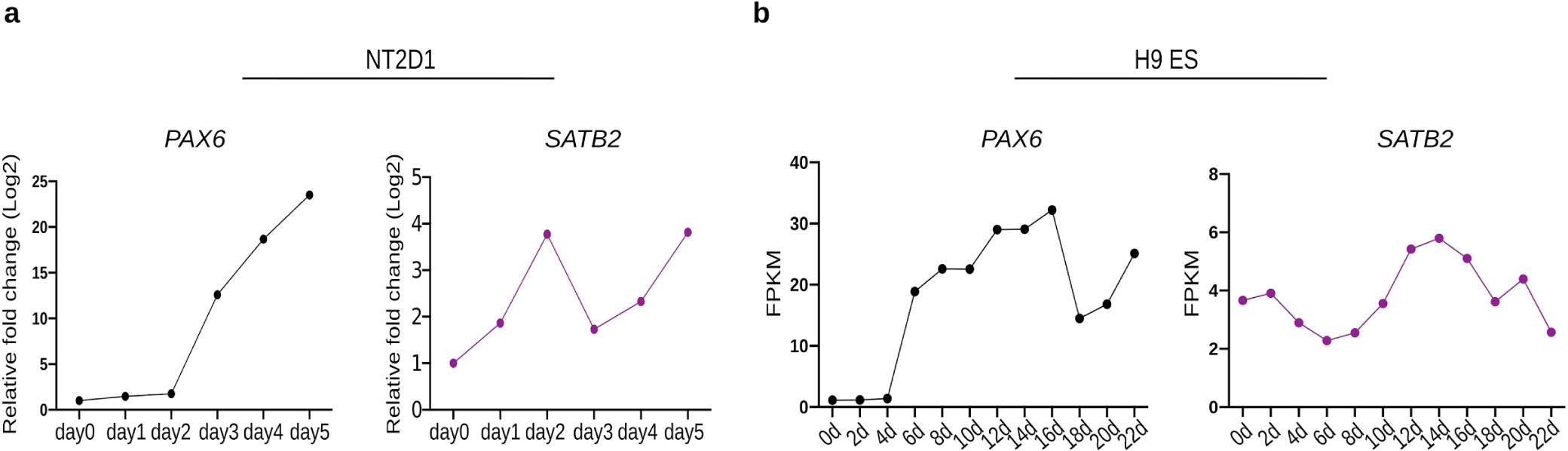
Expression pattern of human *SATB2* during differentiation to neural progenitors and predictive interactors of Satb2 across developmental stages of zebrafish. **a**, Line graph depicting average mRNA expression levels for *PAX6* and *SATB2* in retinoic acid differentiation of NT2D1 cells (n=2). **b**, Line graph representing mRNA expression analysis of *PAX6* and *SATB2* in neural differentiation model in H9 ES cells.

## Notes

### Competing Interest Statement

The authors have declared no competing interest.

